# A multi-layered and dynamic apical extracellular matrix shapes the vulva lumen in *Caenorhabditis elegans*

**DOI:** 10.1101/2020.04.14.041483

**Authors:** Jennifer D. Cohen, Alessandro P. Sparacio, Alexandra C. Belfi, Rachel Forman-Rubinsky, David H. Hall, Hannah M. Maul-Newby, Alison R. Frand, Meera V. Sundaram

## Abstract

Biological tubes must develop and maintain their proper diameter in order to transport materials efficiently. These tubes are molded and protected in part by apical extracellular matrices (aECMs) that line their lumens. Despite their importance, aECMs are difficult to image *in vivo* and therefore poorly understood. The *C. elegans* vulva has been a paradigm for understanding many aspects of organogenesis. Here we describe the vulva luminal matrix, which contains chondroitin proteoglycans, Zona Pellucida (ZP) domain proteins, and other glycoproteins and lipid transporters related to those in mammals. Confocal and transmission electron microscopy revealed, with unprecedented detail, a complex and dynamic aECM. Different matrix factors assemble on the apical surfaces of each vulva cell type, with clear distinctions seen between Ras-dependent (1°) and Notch-dependent (2°) cell types. Genetic perturbations suggest that chondroitin and other aECM factors together generate a structured scaffold that both expands and constricts lumen shape.

## Introduction

During tubulogenesis, lumen formation and expansion generally occur in the context of fluid influx and/or apical extracellular matrix (aECM) secretion (reviewed by (Luschnig & Uv, 2014; Navis & Nelson, 2016)). Tubular epithelia drive water into the lumen by establishing ionic and osmotic gradients using various ion pumps and channels; the resulting hydrostatic pressure can stimulate lumen enlargement (Bagnat, Cheung, Mostov, & Stainier, 2007; Dong, Deng, & Jiang, 2011; Khan et al., 2013; Kolotuev, Hyenne, Schwab, Rodriguez, & Labouesse, 2013; Navis, Marjoram, & Bagnat, 2013). But ions and water are not the only molecules being secreted into nascent lumens; proteoglycans, lipids, mucins, zona pellucida (ZP) domain proteins, and/or other matrix factors are also present and can contribute to lumen shaping (Devine et al., 2005; Gill et al., 2016; Hwang, Olson, Esko, & Horvitz, 2003; Jazwinska, Ribeiro, & Affolter, 2003; Lane, Koehl, Wilt, & Keller, 1993; Rosa, Metzstein, & Ghabrial, 2018; Strilić et al., 2009; Tonning et al., 2005). These aECM factors may act like sponges to bind and organize water molecules and generate outward pushing forces (Lane et al., 1993; Syed et al., 2012), or they may assemble into fibrils or other specialized structures to exert more localized pushing or pulling forces on tube membranes (Andrew & Ewald, 2010; Linde-Medina & Marcucio, 2018; Luschnig & Uv, 2014; Plaza, Chanut-Delalande, Fernandes, Wassarman, & Payre, 2010). aECMs may also bind and present or sequester various signaling molecules that impact cell identity or behavior (Judge & Dietz, 2005; Perrimon & Bernfield, 2000). aECMs of varying types are present in all tubular epithelia; examples in mammals include the vascular glycocalyx, lung surfactant and the mucin-rich linings of the gastrointestinal tract and upper airway (Bernhard, 2016; Johansson, Sjovall, & Hansson, 2013; Webster & Tarran, 2018). However, such aECMs generally appear translucent by light microscopy and are easily destroyed by standard chemical fixation protocols, and thus the organizational structures and lumen-shaping mechanisms of most luminal matrices remain poorly understood.

Vulva development in the nematode *Caenorhabditis elegans* has been a paradigm for understanding many aspects of cell fate specification and organogenesis (Schindler & Sherwood, 2013; Schmid & Hajnal, 2015). The vulva tube consists of twenty-two cells of seven different cell types, organized as seven stacked toroids (vulA, vulB1, vulB2, vulC, vulD, vulE and vulF) (Sharma-Kishore, White, Southgate, & Podbilewicz, 1999) (Figure 1). In adult hermaphrodites, the vulva connects to the uterus and serves as a passageway to allow sperm entry and the release of fertilized eggs. In the forty+ years since vulva cell lineages were first described (Sulston & Horvitz, 1977), much has been learned about how different vulva cell fates are specified and how they arrange to form the tube structure. It is known that the glycosoaminoglycan (GAG) chondroitin promotes initial expansion of the vulva lumen during morphogenesis (Hwang, Olson, Esko, et al., 2003), the lumen changes shape and eventually narrows, and then later, in the adult, collagenous cuticle lines the functional vulva tube (Page & Johnstone, 2007; Sulston & Horvitz, 1977). However, the specific contents, organization, and morphogenetic roles of the luminal matrix within the developing vulva have remained, for the most part, uncharacterized.

**Figure 1.**
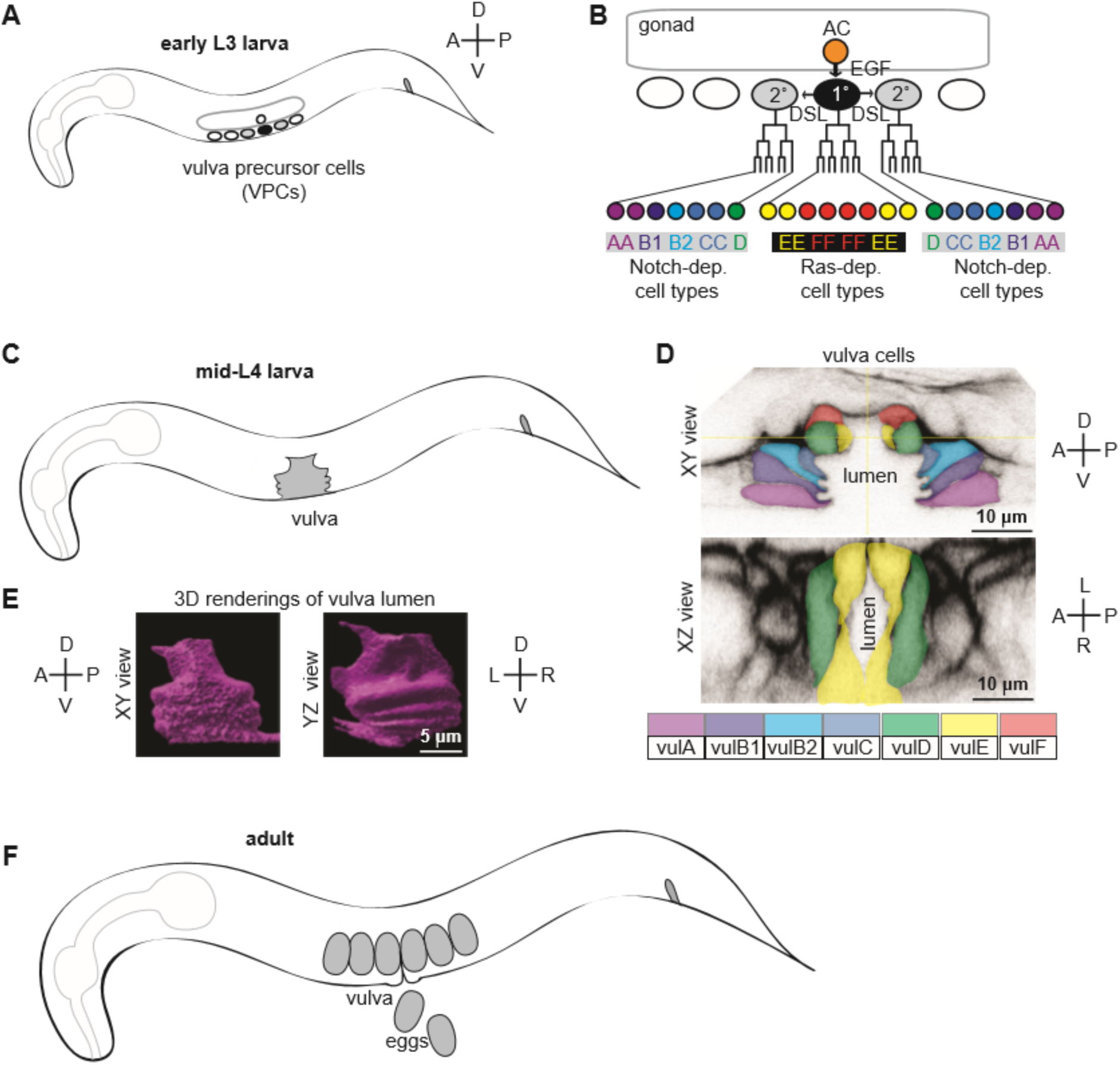
Introduction to vulva development. A) Cartoon of early L3 larva, indicating the six Vulva Precursor Cells (VPCs) beneath the somatic gonad. B) Vulva lineages and cell types. An EGF-like signal from the gonadal anchor cell (AC) induces the primary (1°) cell fate in the nearest VPC (black), which then expresses DSL ligands to induce the secondary (2°) cell fate in the adjacent VPCs (grey). The 1° and 2° VPCs divide to generate a total of twenty-two descendants of seven different cell types. C) Cartoon of mid L4 larva, showing the vulva lumen. D) L4.4 stage vulva cells visualized with the membrane marker MIG-2::GFP (*muIs28*). The twenty-two vulval cells are organized into seven stacked rings (Sharma-Kishore et al., 1999). In the standard lateral or sagittal view, anterior is to the left and ventral is down. An orthogonal XZ view shows the oblong shape of the lumen. E) 3D rendering of the L4.4 vulva lumen generated with Imaris software (BitPlane, Zurich Switzerland), based on imaging of the matrix factor FBN-1 (see Figure 3). The YZ view at right is comparable to the transverse views of the vulva seen by TEM in Figures 4 and 8C (but note that regions deeper in the Z plane are poorly resolved here). F) The adult vulva is a slit-like and cuticle-lined passageway through which eggs are laid.

Here we show that a spatially and temporally dynamic aECM assembles and disassembles within the vulva lumen during morphogenesis. This aECM shares components with the glycocalyx-like sheath or pre-cuticle matrix that coats other apical surfaces in *C. elegans* prior to each round of cuticle secretion (Forman-Rubinsky, Cohen, & Sundaram, 2017; Gill et al., 2016; Katz, Maybrun, Maul-Newby, & Frand, 2018; Kelley et al., 2015; Labouesse, 2012; Lazetic & Fay, 2017; Mancuso et al., 2012; Priess & Hirsh, 1986; Vuong-Brender, Suman, & Labouesse, 2017). It contains both fibrillar and gel-like or granular components, and also extracellular vesicles, as observed at the ultrastructural level. Different combinations of matrix factors assemble on the apical surfaces of each of the seven different vulva cell types, with particularly clear distinctions seen between Ras-dependent (1°) and Notch-dependent (2°) cell types. Genetic perturbation experiments suggest that chondroitin and other aECM factors together generate a structured scaffold that has both lumen expanding and lumen constraining roles.

## Results

### Background: vulva tube formation

Specification and generation of the seven vulva cell types occurs during the L2 and L3 larval stages, while toroid formation and other aspects of tube morphogenesis occur during the L4 stage (Figures 1 and 2). The twenty-two cells of the vulva derive from three of six total possible vulva precursor cells (VPCs), named P3.p-P8.p. Signaling by the Epidermal Growth Factor Receptor (EGFR)-Ras-ERK and Notch pathways specifies one central 1° and two flanking 2° VPC fates (Figure 1B) (Schmid & Hajnal, 2015; Sternberg & Horvitz, 1989). First, an EGF-like signal from the gonadal anchor cell (AC) induces P6.p to adopt the 1° VPC fate, and that cell then expresses Delta/Serrate/LAG-2 (DSL)-like ligands to induce its neighbors, P5.p and P7.p, to adopt the 2° cell fate. The 1° VPC (P6.p) divides to generate eight descendants: 4 vulF and 4 vulE cells. The 2° VPCs (P5.p and P7.p) divide to generate seven descendants each: 1 vulD, 2 vulC, 1 vulB2, 1 vulB1 and 2 vulA cells. As they divide, primary descendants produce an unknown cue that promotes basement membrane invasion by the gonadal AC, which is the first step in forming a vulva-uterine connection (Sherwood & Sternberg, 2003). 1° descendants also begin to detach from the underlying epidermal cuticle and move dorsally. Upon completion of vulva cell divisions, 2° descendants migrate inward in a Rac and Rho-dependent manner, and push the more central 2° descendants and the 1° descendants further dorsally to generate the vulva invagination seen at early L4 (Farooqui et al., 2012; Kishore & Sundaram, 2002; Sharma-Kishore et al., 1999; Shemer, Kishore, & Podbilewicz, 2000). As cells of the same type meet, they eventually fuse to form the seven vulva toroids (Sharma-Kishore et al., 1999).

**Figure 2.**
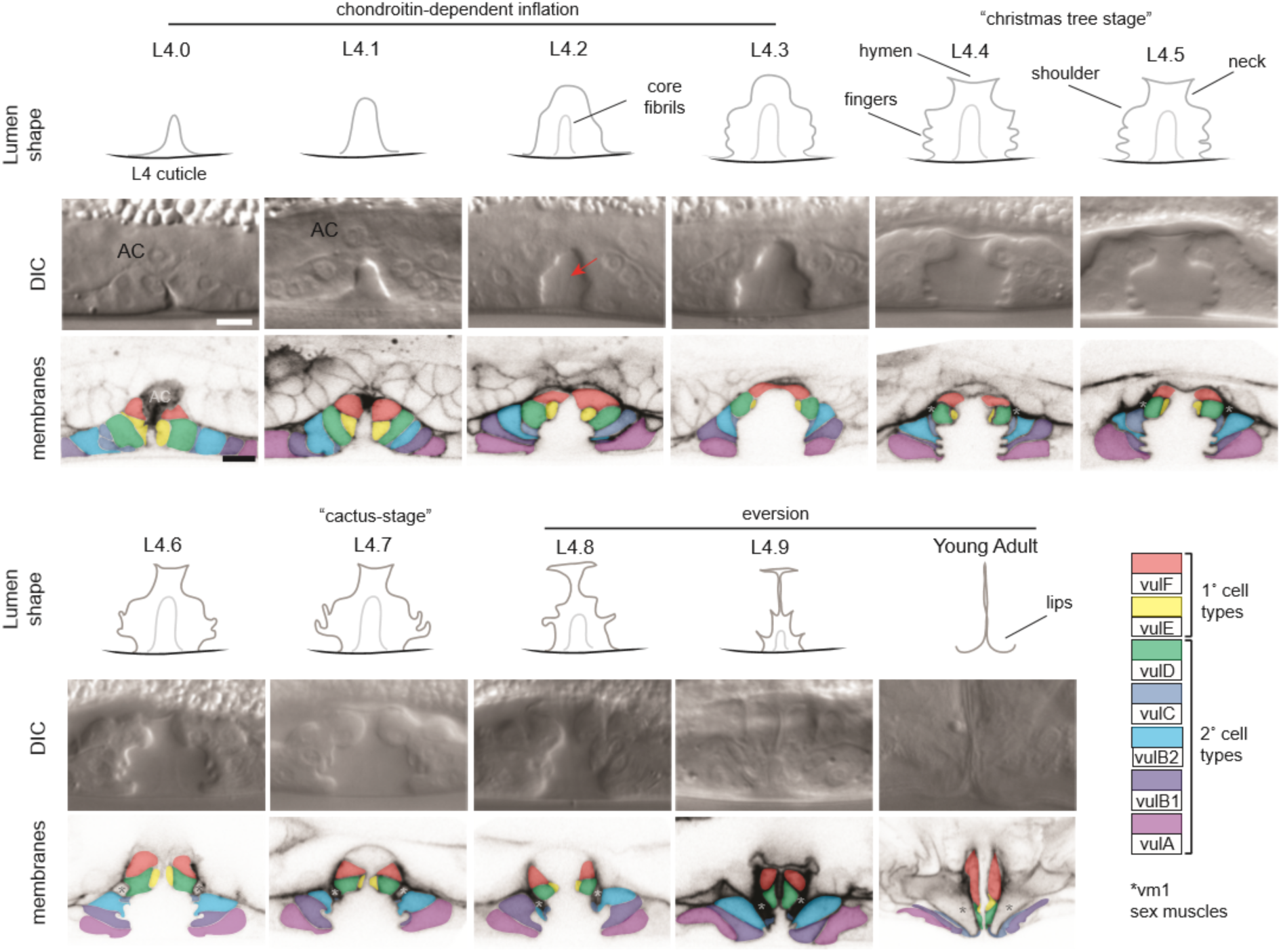
Cell shape changes during vulva morphogenesis. Sagittal views of the central vulva lumen. Top rows show cartoons of lumen shape for each L4 sub-stage, as defined by Mok et al (Mok et al., 2015). Middle rows show corresponding DIC images. Bottom rows show confocal slices of vulva cell membranes marked by MIG-2::GFP (*muIs28*); cells are colored according to the key at right. Luminal core fibrils are faintly visible beginning at L4.3 (red arrow). At mid-L4 (“Christmas-tree stage”; (Seydoux et al., 1993)), the vulF and vulE cells together define the vulva “neck”, the vulD and vulC cells define the vulva “shoulder”, the vulB1 and vulB2 cells define the vulva “fingers”, and the vulA cells make the connection between the vulva cells and the surrounding epidermis. *vm1 sex muscles, which attach to the mature vulva between the vulC and vulD toroids (Sharma-Kishore et al., 1999). Scale bars, 5 microns.

Ten different stages of L4 vulva morphogenesis (L4.0-L4.9) have been distinguished based on changing lumen morphology, as observed by differential interference contrast (DIC) microscopy (Mok, Sternberg, & Inoue, 2015). To visualize cell shapes that correspond to each of these stages, we used the RhoG marker MIG-2::GFP (Honigberg & Kenyon, 2000) to label all vulva cell membranes (Figures 1D, 2). At L4.0, the vulva invagination is very narrow, but it enlarges to approximately 10 microns in diameter by the L4.3 stage. Between L4.3 and L4.4, the uterine lumen and the dorsal-most part of the vulva lumen both expand, and the gonadal AC fuses with the uterine seam (utse), leaving just a thin part of its membrane as a hymen separating the two lumens (Sapir et al., 2007; Sharma-Kishore et al., 1999). The vulB1 and vulB2 cells also develop increasingly concave apical surfaces, creating a “Christmas tree-like” lumen appearance at L4.4-L4.5, and a more “cactus-like” appearance by L4.7. In the final morphogenetic stages, collectively termed eversion, further cell shape changes and rearrangements occur to narrow the lumen and generate the closed lips of the final adult tube structure (Seydoux, Savage, & Greenwald, 1993) (see below). Following eversion, the vulva lumen remains in a closed conformation unless opened by contractions of the sex muscles, which attach to multiple vulva toroids (Sharma-Kishore et al., 1999). During L4 and adult stages, each vulva cell type expresses different combinations of known transcription factors, membrane fusogens, or other molecular markers (Inoue et al., 2004; Inoue et al., 2002; Inoue, Wang, Ririe, Fernandes, & Sternberg, 2005; Mok et al., 2015; Shemer et al., 2000; Sternberg & Horvitz, 1989), but the biological distinctions among the seven cell types are not well understood.

A chondroitin proteoglycan (CPG)-rich luminal matrix is thought to form at the earliest stages of vulva tube morphogenesis, and to swell with water to exert a uniform pushing force for the lumen expansion (Gupta, Hanna-Rose, & Sternberg, 2012; Hwang, Olson, Esko, et al., 2003). Chondroitin antibodies stain the mid-L4 vulva lumen, though in a disorganized fashion that likely reflects matrix destruction by chemical fixatives (Bender, Kirienko, Olson, Esko, & Fay, 2007). Mutants defective in chondroitin biosynthesis have a narrow or “squashed” vulva lumen (Sqv phenotype) (Herman, Hartwieg, & Horvitz, 1999; Hwang, Olson, Esko, et al., 2003). Genetic screens for Sqv mutants identified many components of the chondroitin biosynthesis pathway (Herman et al., 1999; Hwang, Olson, Brown, Esko, & Horvitz, 2003; Hwang, Olson, Esko, et al., 2003), but did not identify any specific chondroitin-modified proteins, suggesting redundant contributions of multiple CPGs. To date, mass spectrometry has identified 24 CPG carrier proteins in *C. elegans*, but it is not yet known which, if any, of these contribute to vulva lumen expansion (Noborn et al., 2018; Olson, Bishop, Yates, Oegema, & Esko, 2006). One of these known CPGs is FBN-1, a fibrillin-related ZP protein that is part of the worm’s embryonic sheath matrix (Kelley et al., 2015; Labouesse, 2012; Noborn et al., 2018; Priess & Hirsh, 1986). Other sheath proteins also have been observed within the vulva (Forman-Rubinsky et al., 2017; Gill et al., 2016), and the adult vulva becomes cuticle-lined in adults (Page & Johnstone, 2007; Sulston & Horvitz, 1977). These observations suggested to us that a sheath-like matrix may exist within the developing vulva lumen.

### Sheath aECM proteins show dynamic patterns of localization during vulva lumen morphogenesis

To examine sheath protein expression and localization in the vulva, we examined Superfolder (Sf) GFP or mCherry-based translational fusions generated by transgenic methods or (in most cases) by CRISPR-Cas9 genome editing of the endogenous loci (Materials and Methods). Six different sheath matrix proteins were assessed (Figure 3A): the ZP proteins FBN-1 (Kelley et al., 2015), LET-653 (Gill et al., 2016) and NOAH-1 (Vuong-Brender et al., 2017), the lipocalin LPR-3 (Forman-Rubinsky et al., 2017), and the extracellular leucine-rich repeat only (eLRRon) proteins LET-4 (Mancuso et al., 2012) and SYM-1 (Davies, Spike, Shaw, & Herman, 1999; Vuong-Brender et al., 2017). We also examined a shortened version of LET-653 containing only the ZP domain (Gill et al., 2016) (Figure 3A). FBN-1, NOAH-1 and LET-4 each have transmembrane domains (but could be cleaved extracellularly), while LET-653, LPR-3 and SYM-1 are secreted proteins. These fusions permitted observations of matrix structure in live L4 animals, without the matrix destruction typically induced by chemical fixation for immunofluorescence. In each case, these fusion proteins were functional as assessed by mutant rescue or strain phenotypes (Materials and Methods), suggesting that their localization closely approximates the endogenous patterns. Each fusion protein initially appeared at around the L4.2 stage, and then exhibited a temporally and spatially distinct pattern over the course of L4 vulva morphogenesis (Figure 3B-D). All six matrix proteins were transient and disappeared by the time that animals reached the adult stage, indicating that they are not components of the mature cuticle, but rather define a distinct matrix type present only during morphogenesis (Forman-Rubinsky et al., 2017; Gill et al., 2016) (and data not shown).

**Figure 3.**
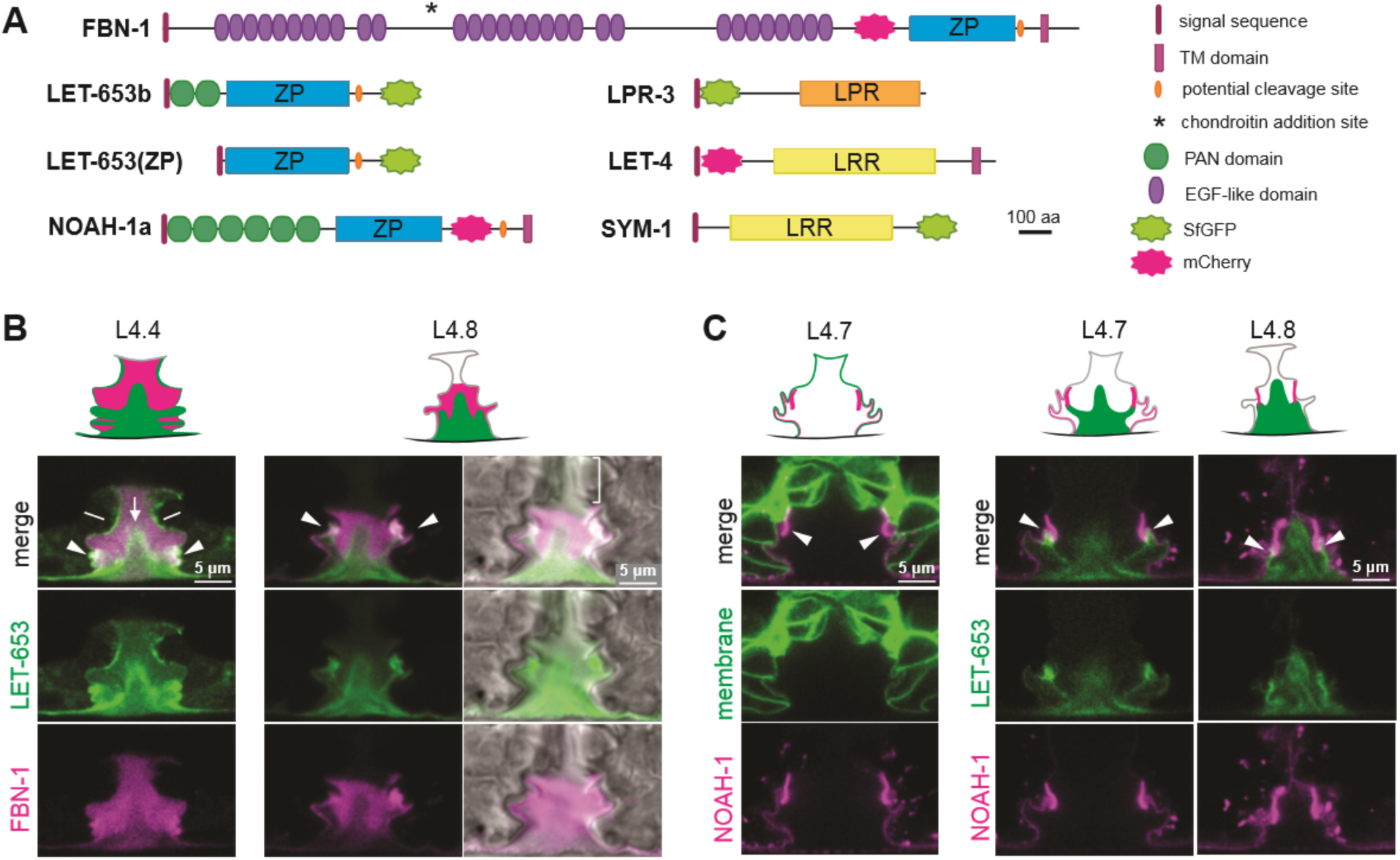
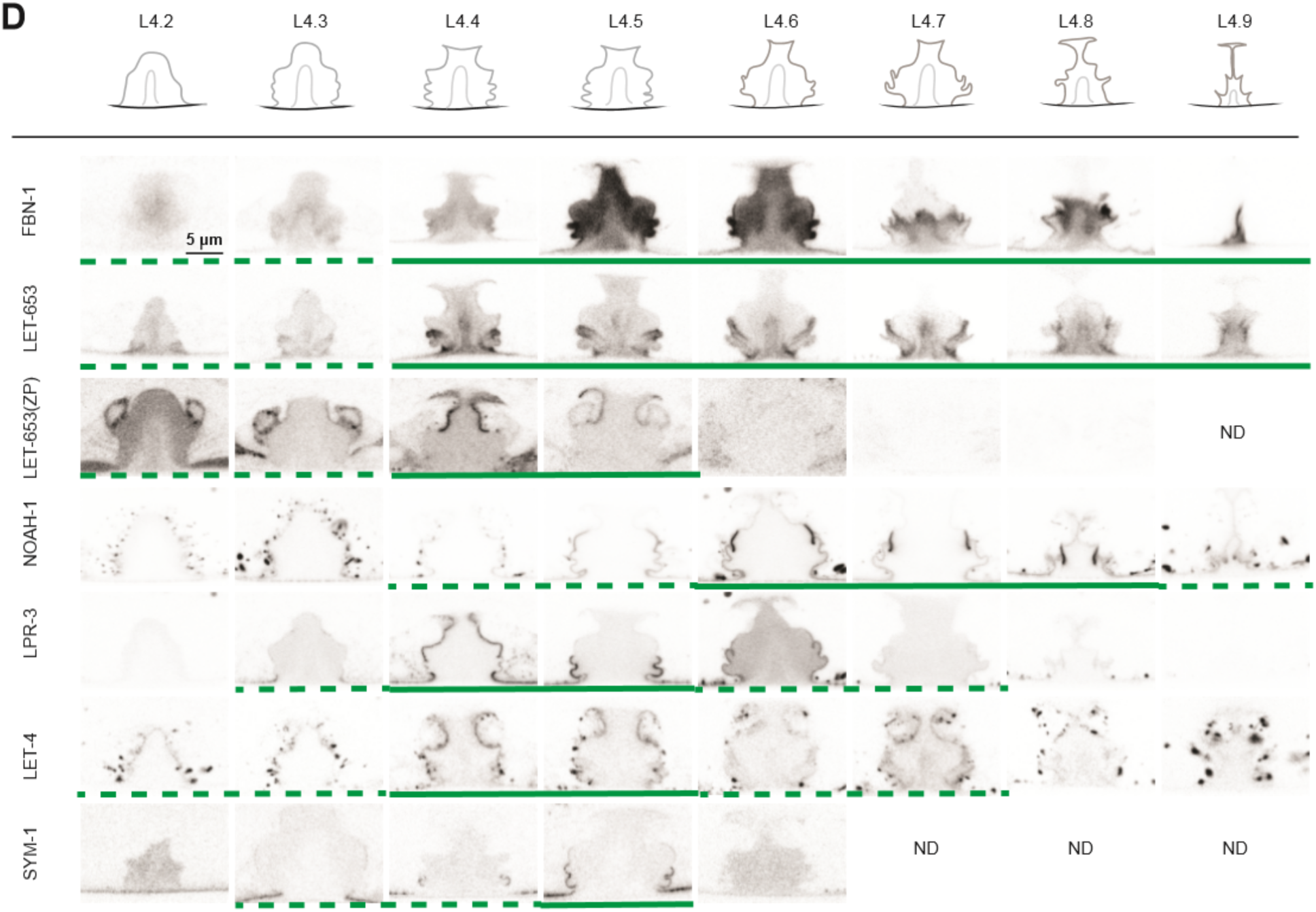
A dynamic aECM fills the vulva lumen during morphogenesis. A) Sheath matrix protein schematics. Genbank accession: FBN-1a (AFN70749.1), LET-653b (CAH60755.1), NOAH-1a (CCD66686.2), LPR-3 (CAA92030.1), LET-4 (AEZ55699.1), SYM-1 (CAB43345). Full-length FBN-1::mCherry and some LET-653 fusions were expressed from transgenes; all others were expressed from the endogenous loci tagged by CRISPR-Cas9 genome editing (see Materials and Methods). B) LET-653::SfGFP (*csIs64*) and FBN-1::mCherry (*aaaIs12*) show complementary luminal patterns. Medial confocal slices. Arrow, core fibrils. Arrowheads, sites of ventrolateral fibril attachment to vulva cells. Lines, membrane-anchored matrix. Bracket indicates loss of FBN-1 from the lumen over 1° cells during vulva eversion. C) NOAH-1::mCherry (*mc68*) labels matrix spikes that connect to LET-653-marked luminal fibrils during vulva eversion. Left column shows overlay with cell membrane marker MIG-2::GFP (*muIs28*). Right columns show overlay with LET-653::SfGFP (*cs262*). D) Timeline of vulva morphogenesis showing dynamic matrix patterns. Each image is a single confocal slice, inverted for clarity. For each fusion, images were collected for at least three animals per stage per strain; most fusions were imaged in multiple different strains to directly compare the different patterns (as in panels B and C). Solid green underlines indicate stages with consistent and peak localization; dashed green underlines indicate stages with more variable or weak localization. Fusions shown are FBN-1::mCherry (*aaaIs12*), LET-653::SfGFP (*cs262*), LET-653(ZP)::SfGFP (*csIs66*), SfGFP::LPR-3 (*cs250*), NOAH-1::mCherry (*mc68*), mCherry::LET-4 (*cs265*), and SYM-1::GFP (*mc85*).

The ZP proteins FBN-1 and LET-653 showed somewhat complementary luminal patterns (Figure 3B,D). Beginning at L4.2, and as previously reported for LET-653(PAN domain) fragments (Gill et al., 2016), LET-653 decorated a core of fibrils in the center of the lumen that rise to the level of the vulD and vulE cells, along with lateral fibrils that connect this core to the vulA, vulB1 and vulB2 cells and to the surrounding epidermis (Figure 3B,D). The central core fibrils could also be detected very weakly by DIC (Figure 2). These fibrils changed appearance during vulva eversion, but remained visible in the most ventral, 2°-cell-derived region through the L4.9 stage. Transiently, at the L4.3-L4.5 stages, LET-653 also weakly marked the apical membranes of most cells. Finally, FBN-1 overlapped with LET-653-marked fibrils near vulB1 and vulB2 surfaces, but otherwise was mainly excluded from the core fibril area and instead filled the more dorsal part of the lumen above the fibrils (Figure 3B,D). During vulva eversion, FBN-1 became excluded from the dorsal-most portions of the lumen lined by 1°-derived cells, such that LET-653 and FBN-1 together appeared to demarcate at least 3 separate luminal zones roughly corresponding to the regions outlined by the vulA/B cells, vulC/D cells, and vulE/F cells (Figure 3B).

The isolated LET-653 ZP domain and the other four sheath matrix proteins marked specific apical membrane-proximal regions in a dynamic manner (Figure 3C,D). LET-653(ZP) specifically labelled just the 1°-derived vulE and vulF cell surfaces at L4.3-L4.5 stages. Previous Fluorescence Recovery After Photobleaching (FRAP) studies showed that, while it is present, this pool of LET-653(ZP) is relatively immobile, consistent with matrix incorporation (Gill et al., 2016). NOAH-1 faintly marked all 2° vulva cell surfaces at L4.4-L4.5, but then became increasingly concentrated on vulC and vulD. During vulva eversion, NOAH-1 prominently marked matrix spikes that protruded from vulC into the lumen, and these spikes attached to LET-653-marked fibrils after those fibrils detached from vulB1 and vulB2 surfaces (Figure 3C,D). These NOAH-1-LET-653 connections then persisted as the lumen narrowed. LPR-3 briefly marked all vulva cell apical surfaces at early L4.4, but then became restricted to 2° cells and then specifically to vulB1 and vulB2 before largely disappearing by L4.6-L4.7 (Figure 3D). The departure of LPR-3 from vulC and vulD coincided with the increasingly strong presence of NOAH-1 there. The transmembrane eLRRon protein LET-4 marked all vulval apical membranes during the late L4.2-L4.7 period, and thereafter appeared intracellular (Figure 3D). Finally, the secreted eLRRon protein SYM-1 showed the most limited pattern, labelling vulB1 and vulB2 for just a brief period at L4.4-L4.5 (Figure 3D). Together, these data reveal that different combinations of sheath matrix factors assemble on the luminal surface of each vulva cell type. Furthermore, the precise timing of each factor’s appearance and disappearance points to highly regulated mechanisms for matrix assembly and remodeling.

### Ultrastructural features of the luminal matrix differ between 1° and 2°-derived vulva regions

Prior transmission electron microscopy (TEM) studies of the vulva (Gill et al., 2016; Herman et al., 1999) used chemical fixation methods that poorly preserved the luminal matrix and did not capture the complex luminal structures observed in the live imaging above. To obtain a clearer view of matrix ultrastructure, we turned to high pressure freezing (HPF) and freeze substitution (Hall, Hartwieg, & Nguyen, 2012). This method achieved much better matrix preservation and revealed many matrix layers and fibrils that we could correlate with those observed by light microscopy. Serial thin sections were collected transverse or length-wise to the body axis to obtain a three-dimensional view. Images of mid-L4 (L4.4-L4.5) and late-L4 (L4-8-L4.9) stage vulvas are shown in Figures 4, 5 and S1, S2. A striking feature of both stages is the difference in matrix organization in the dorsal (1°-derived) vs. ventral (2°-derived) portions of the lumen.

**Figure 4.**
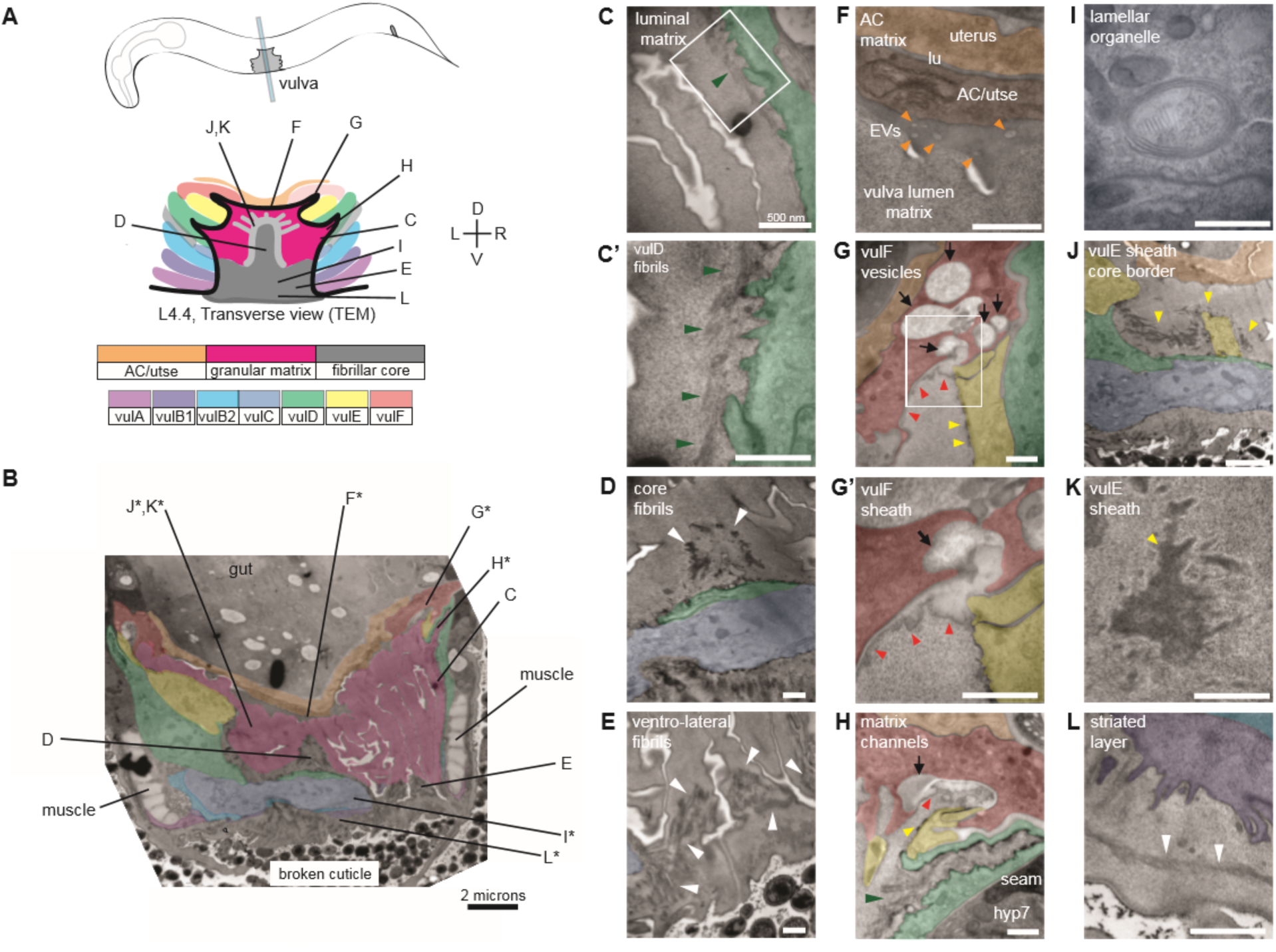
Ultrastructural features of the mid-L4 vulva aECM. A) Transverse serial thin sections of an N2 L4.4 stage animal were analyzed by TEM. The cartoon depicts the vulva lumen in this orientation (see also Figure 1E), and lines indicate the relative locations of different panel images. B) Whole vulva view, L4.4-L4.5 stage. This thin section captures a portion of the lumen and the cell borders. Vulva cells are pseudo-colored according to the key shown in A. The space-filling granular matrix is colored pink to match cartoon. Lines indicate the relative locations of different panel images, and asterisks indicate that the panel shows a region from a different thin section of the same animal. The ventral cuticle has broken during specimen processing and oval objects surrounding the specimen are *E. coli* bacteria. C and C’) A rough granular matrix fills the dorsal lumen. Inset in C’ shows fine fibrils embedded within this matrix (black arrowheads), adjacent to the protrusive surface of vulD. D) Core fibrils (white arrowheads) rise above vulC and vulD, to the level of vulE (which is not present in this section). E) Ventro-lateral fibrils (white arrowheads) and the ventral edge of the luminal matrix, which has pulled away from the broken cuticle. F) A darkly staining fine-grained matrix lines the AC/utse and contains numerous EVs (orange arrowheads). G and G’) vulF cells contain large secretory vesicles (arrows) that empty their contents into the dorsal lumen channels and populate the membrane-anchored matrices that line vulF and vulE (red and yellow arrowheads, respectively). G and H) vulD forms smaller channels that are densely populated with fine fibrils (black arrowheads). I) Lamellar organelle in vulB cell. J) Lateral view of the matrix lining vulE surfaces. K) Closeup of vulE matrix. L) Examples of the very protrusive cell surfaces that interface with the core and ventro-lateral fibrils. L also shows the ventral-most border of the luminal matrix, which contains a striated layer (white arrowheads) similar to that seen in epidermal cuticle (Page & Johnstone, 2007). All scale bars are 500 nm unless otherwise indicated. See Figure S1 for uncolored versions of all images.

**Figure 5.**
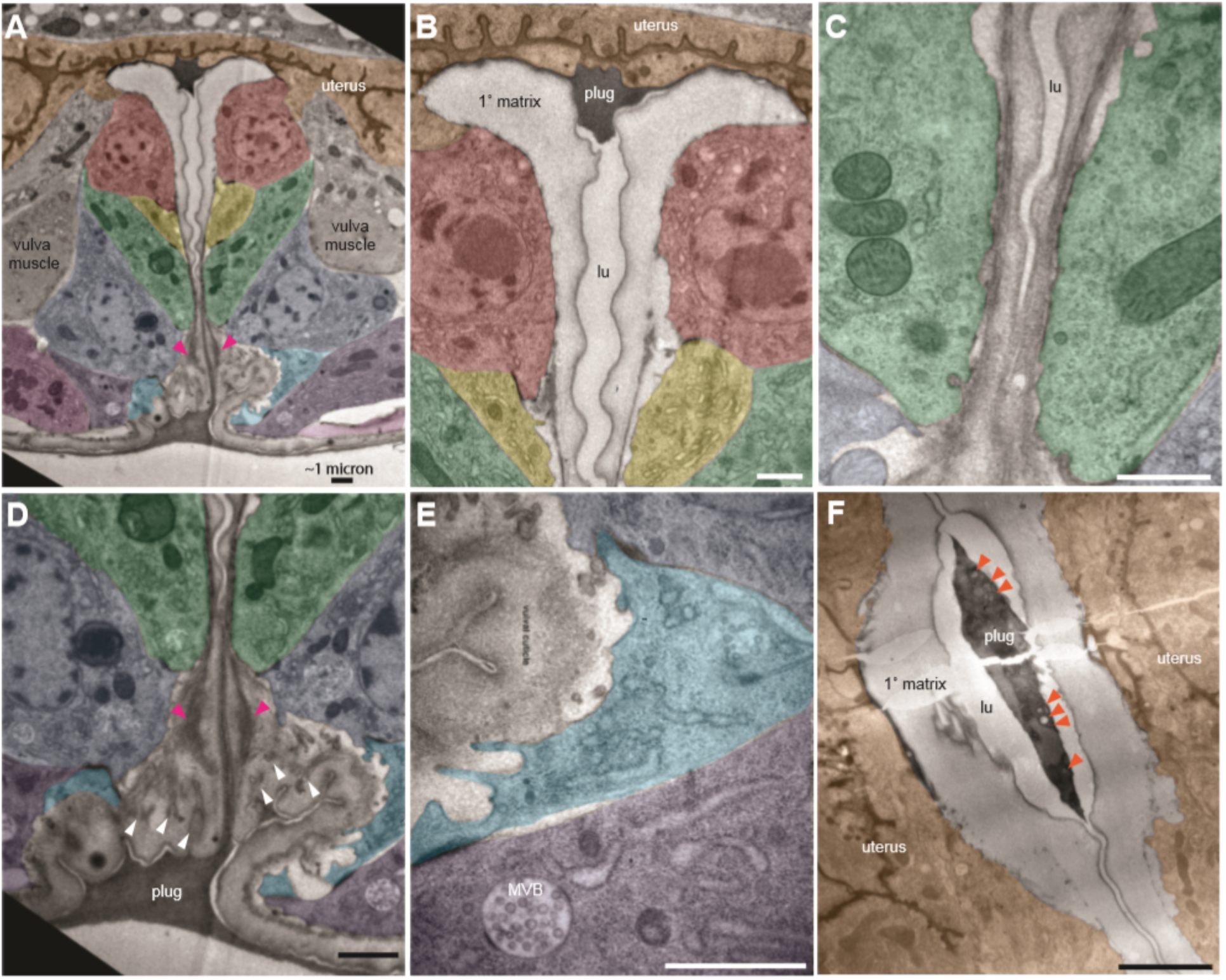
Ultrastructural features of the late L4 vulva aECM. A) Longitudinal slice through the vulva of an N2 L4.8-L4.9 stage animal, with orientation similar to that in confocal images. Vulva and uterine cells are pseudo-colored as in Figure 4B. Pink arrowheads indicate matrix spikes as observed with NOAH-1::mCherry (see Figure 3C). B-E) Higher magnification views of the specimen in panel A. B) Primary vulva cells are covered in a thick membrane-anchored matrix, and the dorsal-most edge of the lumen is filled with a plug of darkly-staining material. C) The membrane-anchored matrix continues over vulD and vulC, but becomes filled with dense fibrils. D) Matrix spikes (pink arrowheads) extend from vulC/D into a cuticle-like matrix below. Various other fibrils (white arrowheads) are present within this ventral matrix. E) Multi-layered nascent cuticle over vulB2. Note protrusive surface of vulB2 and multi-vesicular body (MVB) within vulB1. Many MVBs are present in vulva cells at this stage. F) Dorsoventral slice through uterine cells and the primary vulva matrix and plug of a second N2 L4.9 stage animal. Note the numerous EVs (orange arrowheads) present within the plug. All scale bars, 1 micron. See Figure S2 for uncolored versions of all images.

At the mid-L4 stage, a rough granular matrix fills the entire vulva lumen, and embedded within it are a central fibrillar core and numerous thick ventro-lateral fibrillar elements similar to those seen with LET-653::SfGFP (Figure 4B-E). Most of the dorsal lumen surrounding the central core contains only the granular matrix; this corresponds to the region marked by FBN-1::mCherry (Figure 3B) and likely contains additional CPGs. There is no single luminal channel running through the granular matrix, which appears organized into multiple wide strips or flaps, each edged with a more electron-dense border (Figure 4B,C). 1°-derived vulval cells have relatively smooth apical surfaces lined with thin matrices that separate them from the granular matrix, while the 2°-derived cells have more protrusive apical surfaces lined with numerous fibrils that are embedded within the granular matrix (Figure 4C’-E).

At the dorsal apex of the vulva, the AC remnant hymen is lined by a layer of finer-grained, electron-dense matrix that separates it from the vulva luminal matrix (Figure 4F). Within this AC matrix are numerous extracellular vesicles (EVs). The contents and purpose of these EVs are not currently known, but the AC is a source of LIN-3/EGF and other signaling molecules (Hill & Sternberg, 1992; Sherwood & Sternberg, 2003).

The vulF cells contain numerous large (∼200 nm) secretory vesicles that resemble those seen in mammalian goblet cells (Figure 4G,G’,H; (Birchenough, Johansson, Gustafsson, Bergström, & Hansson, 2015)). These vesicles contain globules that resemble mucin packets, along with a few membranous intraluminal vesicles (ILVs). The secretory vesicles appear to be dumping their contents within sequestered luminal channels at the left and right extremes of the lumen, and these contents then expand spherically upon contacting the outside environment. The contents then spread from these luminal channels to form a thin membrane-anchored matrix layer that likely corresponds to the layer marked by LET-653(ZP)::SfGFP (see Figure 3D). The other vulval cells all contain smaller and less numerous examples of other likely secretory vesicles and/or multi-lamellar organelles (Figure 4I).

vulE surfaces are decorated by a flat, gray, mesh-like matrix that drapes down along the top border of the core and ventro-lateral fibrils (Figure 4J,K). This matrix appears as dark membrane-associated patches when vulE is viewed in cross-section (Figure 4G,G’). This matrix may serve as the barrier that excludes FBN-1 from the ventral fibrillar region (see Figure 3B,D).

vulD and vulC surfaces that sit above the ventro-lateral fibrils (and external to the core) are lined with thin fibrils that run in a dorsal-ventral orientation, parallel to the cell membranes (Figure 4C,C’). These fibrils are embedded within the granular luminal matrix rather than forming a separate layer, and they abut numerous small cellular projections. The fibrils are particularly concentrated in narrow (∼0.5 micron) lumen pockets generated by the complex shape of vulD (Figure 4H). These fibrillar regions correspond to those surfaces that become strongly marked by NOAH-1::mCherry (see Figure 3C).

Finally, the ventral-most vulC surfaces, as well as vulB2, vulB1 and vulA, interface with the dense ventro-lateral fibrils, which run both perpendicular to and parallel with the cell membranes (Figure 4D,E). The cell surfaces that interface with these fibrils are extremely protrusive (Figure 4L). At the most ventral edge of the lumen, beneath the core fibrils, matrix layers that resemble those of the epidermal sheath and nascent cuticle interface with the remaining L4 epidermal cuticle (which has broken and pulled away somewhat in this specimen) (Figure 4B,E,L).

The late-L4 stage vulva (Figure 5) retains several of the matrix features seen at the earlier stage, with some notable differences. An AC-like matrix remains as a “plug” at the dorsal apex and still contains EVs (Figure 5A,B,F). A similar-appearing matrix is also present at the ventral opening of the lumen (Figure 5A,D). The 1°-derived vulE and vulF cells and the 2°-derived vulC and vulD cells are now covered with a thick membrane-anchored matrix that somewhat resembles the prior CPG matrix, but with a well-defined, darkly staining border and a single open channel that runs through its center (Figure 5A,B,F). The membrane-anchored matrix over vulC and vulD also contains many dense fibrils that extend down into the matrix below (Figure 5C,D); these likely correspond to the NOAH-1-marked matrix spikes observed by confocal imaging (see Figure 3C). The remaining vulA, vulB1 and vulB2 cells are covered with a more complex, multi-layered matrix that resembles the nascent pre-cuticle on nearby epidermal cells (Figure 5D,E). Various fibrillar structures are embedded within this thick matrix (Figure 5D), possibly corresponding to the core and ventrolateral fibrils seen earlier. Thus, just as seen by confocal light microscopy at this stage (Figure 3B), TEM shows 3 distinct luminal zones corresponding to the regions outlined by the vulA/B cells, vulC/D cells, and vulE/F cells.

### 1° and 2° vulva cell types produce and assemble different matrices

To better understand the differences between the matrix produced and assembled by 1° vs. 2° vulva cell types, we analyzed matrix patterns in *lin-12/Notch* mutants. LIN-12/Notch promotes 2° vs. 1° VPC fates, so loss-of-function [*lin-12(0)*] mutants have only 1° vulva cell types, while gain-of-function [*lin-12(d)*] mutants have only 2° vulva cell types (Greenwald, Sternberg, & Horvitz, 1983; Sternberg & Horvitz, 1989). *lin-12(0)* and *lin-12(d)* mutants both have well-inflated (though mis-shapen) vulva lumens (Figure 6), indicating that relevant CPGs are made by both sets of vulva cell types. Indeed, Herman et al (Herman et al., 1999) previously showed that both types of *lin-12* mutants require the Sqv chondroitin biosynthesis pathway for lumen inflation.

**Figure 6.**
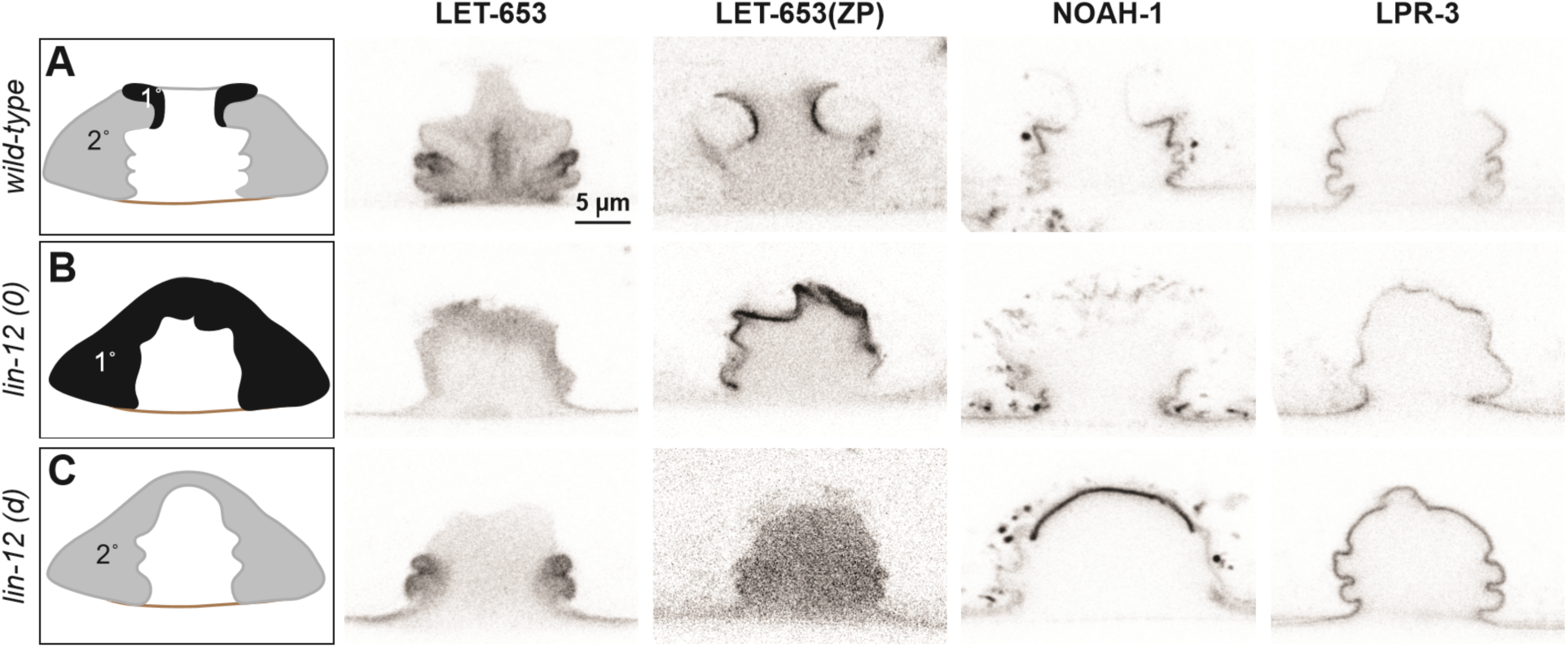
Different vulva cell types produce and assemble different aECMs. A-C) Panels in left column show cartoons of vulva cell types and lumen shape at mid-L4. Remaining columns show single confocal slices through the vulva lumen. Fusions used are LET-653(full-length)::SfGFP (*cs262*), LET-653(ZP)::SfGFP (*csIs66*), NOAH-1::mCherry (*mc68*), SfGFP::LPR-3 (*cs250*). At least n=8 L4s were imaged for each strain. A) In wild type animals, full-length LET-653 predominantly labels the core and ventro-lateral fibrils, LET-653(ZP) labels the membrane-anchored matrix over 1° cells, NOAH-1 labels membrane-anchored matrices over 2° cells (especially vulC and vulD), and LPR-3 transiently labels membrane-anchored matrices over all cells, but then becomes concentrated over 2° cells (see also Figure 3D). B) Loss of 1° cells in *lin-12(0)* (null, *n137n720*) mutants disrupted core fibrils and NOAH-1 localization. Most of the NOAH-1 pattern here is intracellular. C) Loss of 2° cells in *lin-12(d)* (hypermorphic, *n137*) mutants disrupted core fibrils and LET-653(ZP) localization.

Close examination of *lin-12* mutants suggested, however, that the central core fibrils were missing. When LET-653::SfGFP was introduced into *lin-12(0)* mutants, the marked fibrils appeared very meager or absent (Figure 6B). In *lin-12(d)* mutants, no core fibrils were observed, but some ventro-lateral fibrils were still present at the vulB “fingers” (Figure 6C). We conclude that both 1° and 2° cells are required to generate core fibrils, but that 2°-derived cells (most likely vulB1 and vulB2) generate at least some of the ventrolateral fibrils independently.

*lin-12* mutants also showed changes in the localization of membrane-bound matrix factors, as predicted based on the cell fate changes. For example, LET-653(ZP) bound to all vulva apical membranes in *lin-12(0)* mutants, but to none in *lin-12(d)* mutants (Figure 6). Since *let-653* is expressed by all seven vulva cell types (Gill et al., 2016), these data suggest that a 1°-specific partner is required to recruit LET-653(ZP) to the membrane-anchored matrix. In contrast, NOAH-1 was mostly absent from vulva apical membranes in *lin-12(0)* mutants, but strongly bound the dorsal-most apical membranes in *lin-12(d)* mutants (Figure 6). LPR-3 bound some apical membranes in both *lin-12(0)* and *lin-12(d)* mutants, but was more robust in the latter, consistent with its 1° and 2° membrane-binding patterns in wild type (Figure 6). In summary, the results of these experiments indicate that each cell’s identity, rather than its position in the organ, determines what matrix factors assemble on its surface.

## Sheath matrix assembles prior to MUP-4/matrillin expression

The mechanisms by which sheath matrix proteins attach to apical membranes remain unknown, particularly since most either do not have transmembrane domains (Figure 3A), do not require their transmembrane domains for function (Mancuso et al., 2012), or are likely cleaved to release the extracellular domains from their transmembrane domains (Bokhove & Jovine, 2018). However, body cuticle attaches to cell surfaces via hemi-desmosome-like structures (CeHDs) that span the epidermis and link to body muscle (Pásti & Labouesse, 2014). Specifically, the CeHD component VAB-10a (related to spectraplakin) (Bosher et al., 2003) links to the matrillin-like transmembrane proteins MUP-4 (Hong et al., 2001) and MUA-3 (Bercher et al., 2001) to connect the epidermis and cuticle (Suman et al., 2019). Similar complexes have been speculated to anchor the sheath (Vuong-Brender et al., 2017).

To address whether CeHD-linked complexes could anchor sheath matrix in the vulva, we asked where VAB-10a and MUP-4 appear in vulval cells. VAB-10::GFP was present along apical membranes of all vulva cells throughout L4 morphogenesis (Figure 7A). However, MUP-4::GFP appeared only later, at the L4.6-L4.7 stage (Figure 7B). At the L4-adult molt, MUP-4::GFP lined all vulva apical membranes, but it was expressed most prominently in the vulA toroid, which links the vulva to the surrounding epidermis (Figure 7B). Therefore, MUP-4 may help connect the cuticle to CeHDs in vulva cells just as it does in the epidermis. However, different (possibly also CeHD-affiliated) linkers must be involved in recruiting the earlier sheath matrix factors.

**Figure 7.**
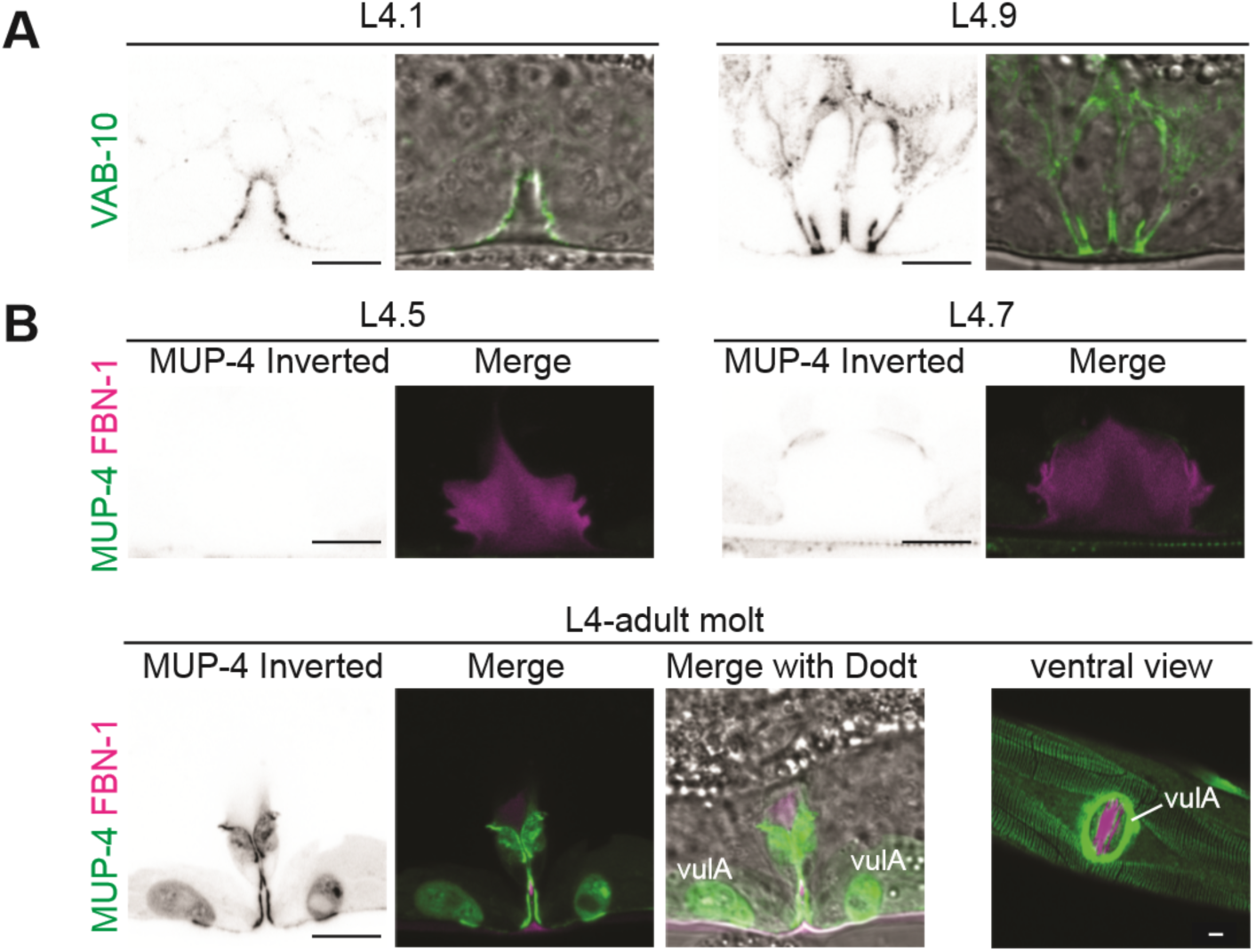
Sheath matrix assembles prior to expression of MUP-4/matrillin. A) VAB-10::GFP (*cas627*) marked all apical membanes in the vulva throughout L4 (including 4/4 L4.1/L4.2 stage animals). B) MUP-4::GFP (*upIs1*) marked apical membranes beginning in late L4 (0/3 L4.4/L4.5, 3/3 L4.6/L4.7, 2/2 L4.9). FBN-1::mCherry (*aaaIs12* or *aaaEx78*) is also shown. At L4.9, MUP-4 expression is particularly strong in the vulA toroid, which surrounds the vulva opening and connects to the surrounding hyp7 epidermis. The remaining FBN-1 matrix connects to vulA at the left and right sides of the lumen, as seen in the ventral view.

### Sheath matrix factors facilitate proper vulva eversion

The complex localization patterns described above suggest important roles for sheath matrix factors in vulva morphogenesis. Of the six sheath factors described here, only *sym-1* mutants are fully viable, while the rest mostly arrest as L1 larvae with excretory tube blockage or other epithelial tissue-shaping defects (Forman-Rubinsky et al., 2017; Gill et al., 2016; Mancuso et al., 2012; Pu, Stone, Burdick, Murray, & Sundaram, 2017; Soulavie, Hall, & Sundaram, 2018; Vuong-Brender et al., 2017). To examine vulva phenotypes in these lethal sheath mutants, we took advantage of rare escapers (for *fbn-1*) or used tissue-specific rescue strains (Materials and Methods) to bypass the earlier requirements (for *let-653* and *let-4*). We were not able to examine *noah-1* or *lpr-3* mutants in this study because of their severe epidermal molting defects (Forman-Rubinsky et al., 2017; Vuong-Brender et al., 2017). All mutants examined had fairly normal vulva lumens at the mid-L4 stage, indicating efficient CPG-dependent lumen inflation (Gill et al., 2016) (Figure 8A). However, in some cases, later stage vulvas appeared mis-shapen or improperly everted (Figure 8A). Specifically, some *let-653* mutants had a prematurely collapsed vulva lumen, and most *fbn-1* mutants had abnormal bulges of the outer vulA, vulB1 and vulB2 cells (Figure 8A). We conclude that sheath factors facilitate later stages of vulva morphogenesis, including vulva eversion.

**Figure 8.**
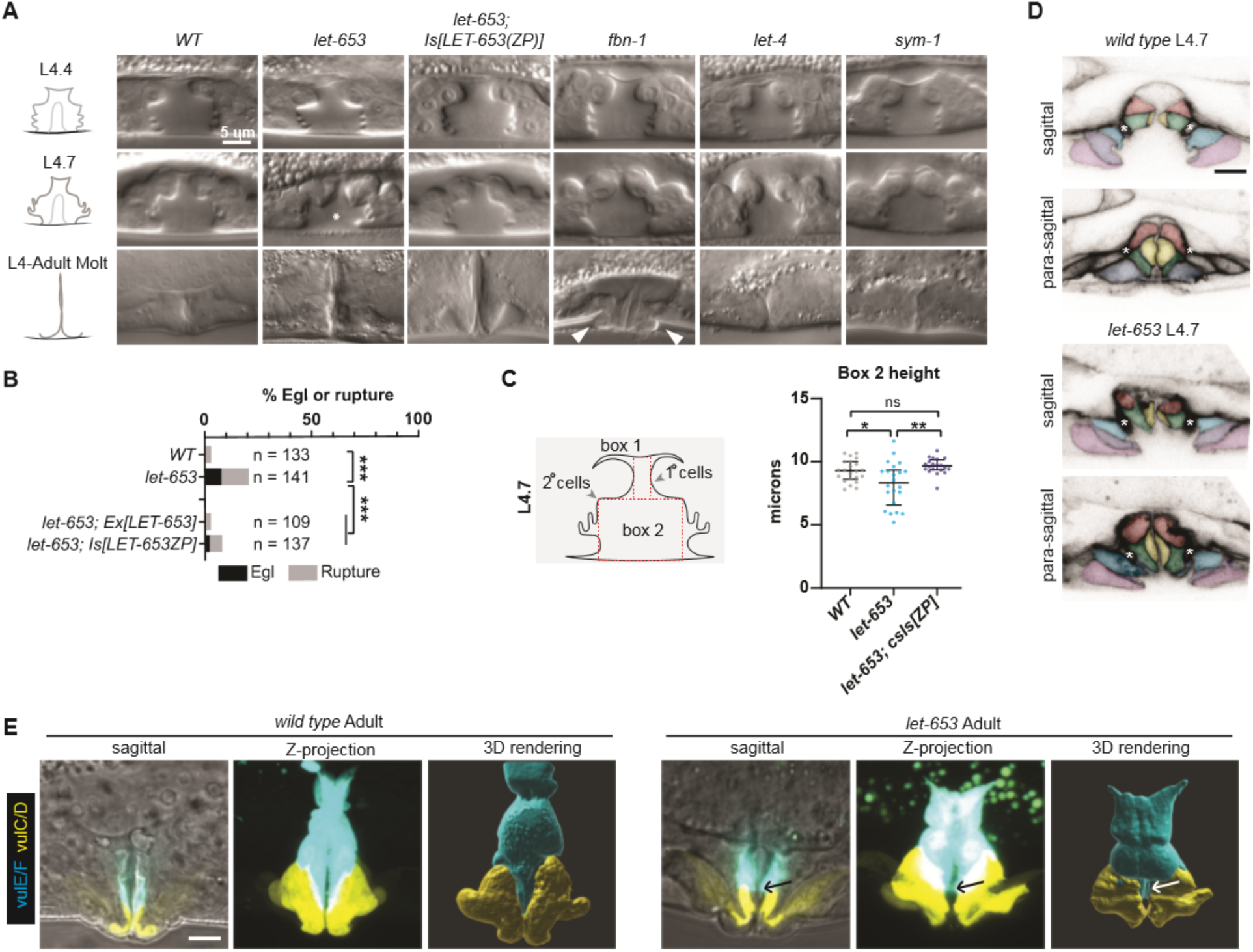
Sheath matrix factors play subtle roles in vulva eversion. A) DIC images of sheath mutant vulvae at L4.4, L4.7 and L4.9-adult molt. At least 40 L4 animals of each genotype were imaged, including at least five each of the three stages shown. Alleles used: *fbn-1(tm290), let-653(cs178)* (strains UP3342 and UP3422), *let-4(mn105), sym-1(mn601)*. Asterisk indicates collapsed lumen morphology in some *let-653* mutants (n=6/22 L4.7, see panel C). Arrowheads indicate abnormal bulges of the vulA and vulB1/B2 cells in *fbn-1* mutants (n=9/13 L4.9). B) A small proportion of *let-653* mutants had progeny that hatched *in utero* (Egl phenotype) or ruptured at the vulva within eight days of reaching adulthood. These phenotypes were rescued by transgenes expressing full-length LET-653 or just the ZP domain. ***p<0.0001, Fisher’s exact test. C) *let-653* mutants have reduced lumen dimensions at the L4.7 stage. To measure lumen dimensions, the largest box possible was drawn within the 1°-generated (Box 1) and 2°-generated (Box 2) lumen spaces, as visualized by DIC. Dimensions for Box 1 did not differ significantly between genotypes (Figure S3), but Box2 height was somewhat reduced. This phenotype was rescued by a transgene expressing the LET-653 ZP domain. *p=0.031, **p=0.001, WT vs. *let-653;Is[ZP]* p=0.085, Mann-Whitney U test. D) *let-653* mutants have irregular vulva cell shapes at the onset of vulva eversion (L4.7 stage). Membranes were visualized with MIG-2::GFP. Asterisks indicate the vm1 sex muscles. Both sagittal and para-sagittal slices from confocal Z-stacks are shown. In wild type, vulva cells are symmetrical across the midline, but in *let-653* mutants, cell shape and position are mismatched (n=3/3; *WT*: n=0/6). E) *let-653* mutants have irregular vulva cell shapes as adults. vulE and vulF were visualized with *daf-6pro::CFP*, and vulC and vulD were visualized with *egl-17pro::CFP* (Mok et al., 2015). Both sagittal slices and Z-projections from confocal Z-stacks are shown, along with three dimensional renderings generated with Imaris (Bitplane) from those Z-stacks. In *let-653* mutants, cell shape and position are variably abnormal (n=3/3; *WT*: 0/3). In the specimen pictured, vulE/F appear less elongated along the dorsal-ventral axis and vulC/D are slightly flattened relative to *WT*. Arrows indicate the transition zone between vulE/F and vulC/D, where increased overlap of the different cell types occurs in the mutant.

To better understand these phenotypes, we focused on *let-653* mutants. As adults, most *let-653* mutants were able to lay eggs; however, a small proportion of mutants ruptured at the vulva and/or were Egg-laying defective (Egl) (Figure 8B). Measurements of the 1°- and 2°-derived portions of the vulva lumen confirmed normal dimensions at the L4.4 stage, but slightly reduced dimensions at the onset of eversion at the L4.7 stage (Figure 8C, Figure S3). A transgene expressing just the LET-653(ZP) domain reversed the L4.7 defects, and led to overexpansion of the lumen at L4.4 (Figure 8C, Figure S3), consistent with our prior report (Gill et al., 2016) that this domain has lumen-expanding properties.

Vulva eversion has been described as the vulva “turning inside out” (Seydoux et al., 1993; Sharma-Kishore et al., 1999), but the specific cellular events involved have never been reported. To visualize cell positions and shapes during eversion, we used the RhoG marker MIG-2::GFP to label all vulva cell membranes (Figures 2, 8D), and the *daf-6pro::CFP* and *egl-17pro::YFP* marker combination (Mok et al., 2015) to label the vulE/F and vulC/D cells specifically (Figure 8E). During wild-type eversion, all four of these cells rotate and elongate in the dorsal-ventral axis to partly occlude the luminal space. vulC extends a narrow NOAH-1 matrix spike into the fibrillar core matrix, whose spokes appear to fold like those of an umbrella as the lumen narrows (Figure 3C). Subsequently, vulA, vulB1 and vulB2 also tilt ventrally, while vulE reaches dorsally to connect to the seam epidermis. By adulthood, vulE and vulF enclose the bulk of the lumen, vulC and vulD form the vulva lips, and the vulA, vulB1 and vulB2 cells are excluded from the lumen, and instead form the epidermis surrounding the vulva opening (Figures 2, 8E).

In *let-653* mutants, vulva eversion occurred in a more irregular manner (Figure 8). The anterior and posterior halves of the vulva showed various asymmetries by the onset of eversion at the L4.7 stage (Figure 8D), and cell shapes were variably abnormal in adults (Figure 8E).

### LET-653 is required to establish well-demarcated dorsal vs ventral matrix domains

To test how LET-653 affects the organization of the vulva matrix, we first assessed the status of other matrix factors in the *let-653* mutant background. *let-653* mutants still assembled some type of central core structure, as seen by DIC and by the exclusion of FBN-1 from this region (Figure 9A). As in WT, FBN-1 departed from the dorsal-most lumen during eversion (Figure 9A) and LPR-3 and NOAH-1 still appeared at their proper locations (Figure 9B). However, the normally precise sequence of LPR-3 clearance was disrupted, such that LPR-3 remained on vulC and vulD apical surfaces longer than normal, and overlapped significantly with NOAH-1 there (Figure 9B).

**Figure 9.**
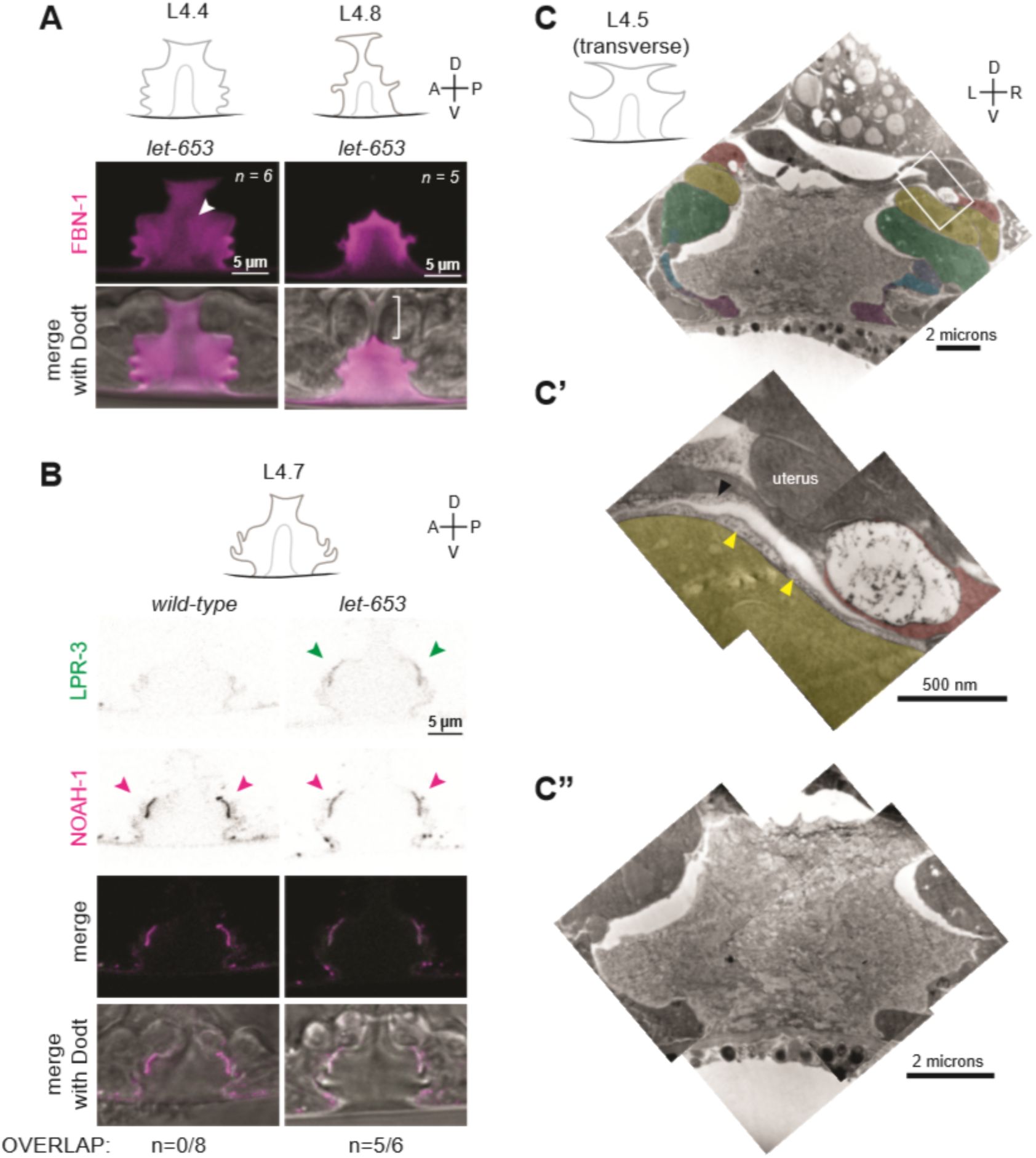
The ZP protein LET-653 is required for proper organization and remodeling of the sheath and luminal matrices. A) *let-653(cs178)* mutants showed relatively normal patterns of FBN-1::mCherry (*aaaIs12*) localization. Arrowhead indicates exclusion of FBN-1 from the core fibril region. Bracket indicates exclusion of FBN-1 from the 1° lumen during eversion. B) *let-653(cs178)* mutants showed normal recruitment of SfGFP::LPR-3 *(cs250)* and NOAH-1::mCherry (*mc68*) to 2° surfaces, but abnormally delayed clearance of SfGFP::LPR-3 from vulC/D. p = 0.0445, Fisher’s exact test. C-C”) Transverse slices of a *let-653(cs178)* mutant at mid-L4 (∼L4.5) stage were analyzed by serial section TEM. Vulva cells are colored as in Figure 4B. C) The overall shape of the vulva appears normal. Box indicates region magnified in C’. C’) An abnormal secretory vesicle in vulF is filled with dark aggregates that match those present in the membrane-anchored matrix over vulF and vulE. C”) Magnified view of the luminal matrix, which contains an ill-defined and patchy core structure.

TEM of a mid-L4 *let-653* mutant revealed more dramatic matrix abnormalities (Figure 9C-C”). Some vulF secretory vesicles contained very disorganized, dark aggregates, and the membrane-anchored matrix over vulF and vulE appeared filled with such aggregates (Figure 9C’). The luminal matrix also contained many aggregates and lacked clearly defined core or ventro-lateral fibrils. Instead, the central core region appeared very patchy, a variety of thin fibrils criss-crossed the entire lumen, and many fibrils accumulated at the dorsal apex of the vulva, along the AC and uterine seam (Figure 9C”). Thus, *let-653* is required to set up an organized core fibril structure and many of the major dorsal vs. ventral differences in the organization of the luminal matrix.

### Chondroitin is not required to assemble a sheath matrix

Thus far, the data indicate that chondroitin plays an early role in vulva lumen inflation, whereas sheath matrix factors play later roles in morphogenesis and eversion. However, at least one sheath factor, FBN-1, is a CPG (Noborn et al., 2018), and other sheath factors appear to be embedded within the CPG matrix (Figure 4), suggesting some coordination between the two processes.

To ask if chondroitin affects sheath matrix assembly, we examined sheath reporters in *sqv-5* and *mig-22* mutants (Figure 10). *sqv-5* and *mig-22* encode chondroitin sulfate synthases related to human CHSY1 and CHSY2 (Hwang, Olson, Esko, et al., 2003; Suzuki, Toyoda, Sano, & Nishiwaki, 2006), which promote chondroitin biosynthesis and polymerization (Izumikawa et al., 2004). *sqv-5* null mutants did still assemble LET-653(ZP) on their 1° surfaces (Figure 10A). However, these null mutants have a very narrow vulva lumen, which made it difficult to stage L4 animals accurately and assess the localization of full length LET-653 or other dynamic luminal factors. Therefore, we turned to *mig-22* hypomorphic (rf=reduced function) mutants, which have a slightly less severe vulva phenotype.

**Figure 10.**
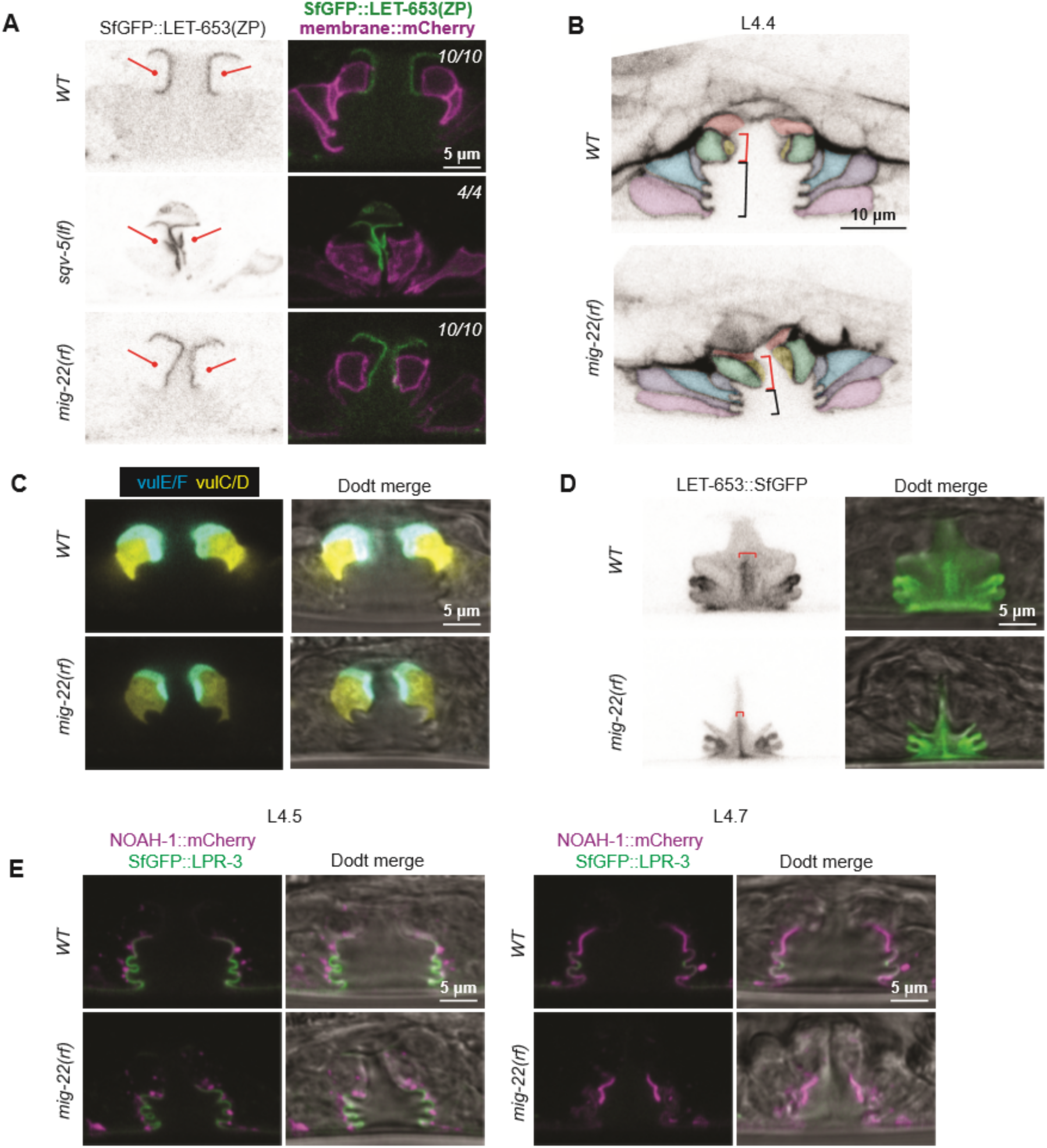
Chondroitin is not required for sheath assembly. A) Chondroitin mutants showed normal recruitment of LET-653(ZP)::SfGFP (*csIs66*) to the membrane-anchored matrix over 1° cells. Alleles used were *sqv-5(n3611)* and *mig-22(k141rf)*. B) WT vs. *mig-22(k141rf)* mutants (L4.4 stage) with membrane marker MIG-2::GFP (*muIs28*). The vulD cells were taller and narrower in mutants compared to wild type, while the vulA, vulB1 and vulB2 cells had shorter apical surfaces (n=15, see also Figure 10B). C) WT vs. *mig-2(k141rf)* mutants (L4.5 stage) with vulE/F marker *daf-6pro::CFP* and vulC/D marker *egl-17pro::YFP* (n=3). D) *mig-22(k141rf)* mutants assembled a well-organized, but narrow, core fibril structure, as seen with LET-653::SfGFP (*cs262*) (n=8). E) *mig-2(k141rf)* mutants showed normal localization of LPR-3::SfGFP and NOAH-1::mCherry to apical surfaces. (n=10)

In *mig-22(rf)* mutants at mid-L4 stage, both the 1°- and 2°-derived parts of the lumen were narrower than in wild-type, and the vulD cells were pre-maturely elongated along the dorsal-ventral axis as previously described for *sqv-3* mutants (Herman et al., 1999) (Figures 10B,C and 11A,B). The vulE and vulF cells appeared relatively normal in shape, but the vulva “neck” region height was greater than wild-type due to the changes in vulD. In contrast, the vulA, vulB1 and vulB2 cells had shorter apical domains (and were instead elongated along the anterior-posterior axis). Thus chondroitin has very different effects on the shape of each cell type.

**Figure 11.**
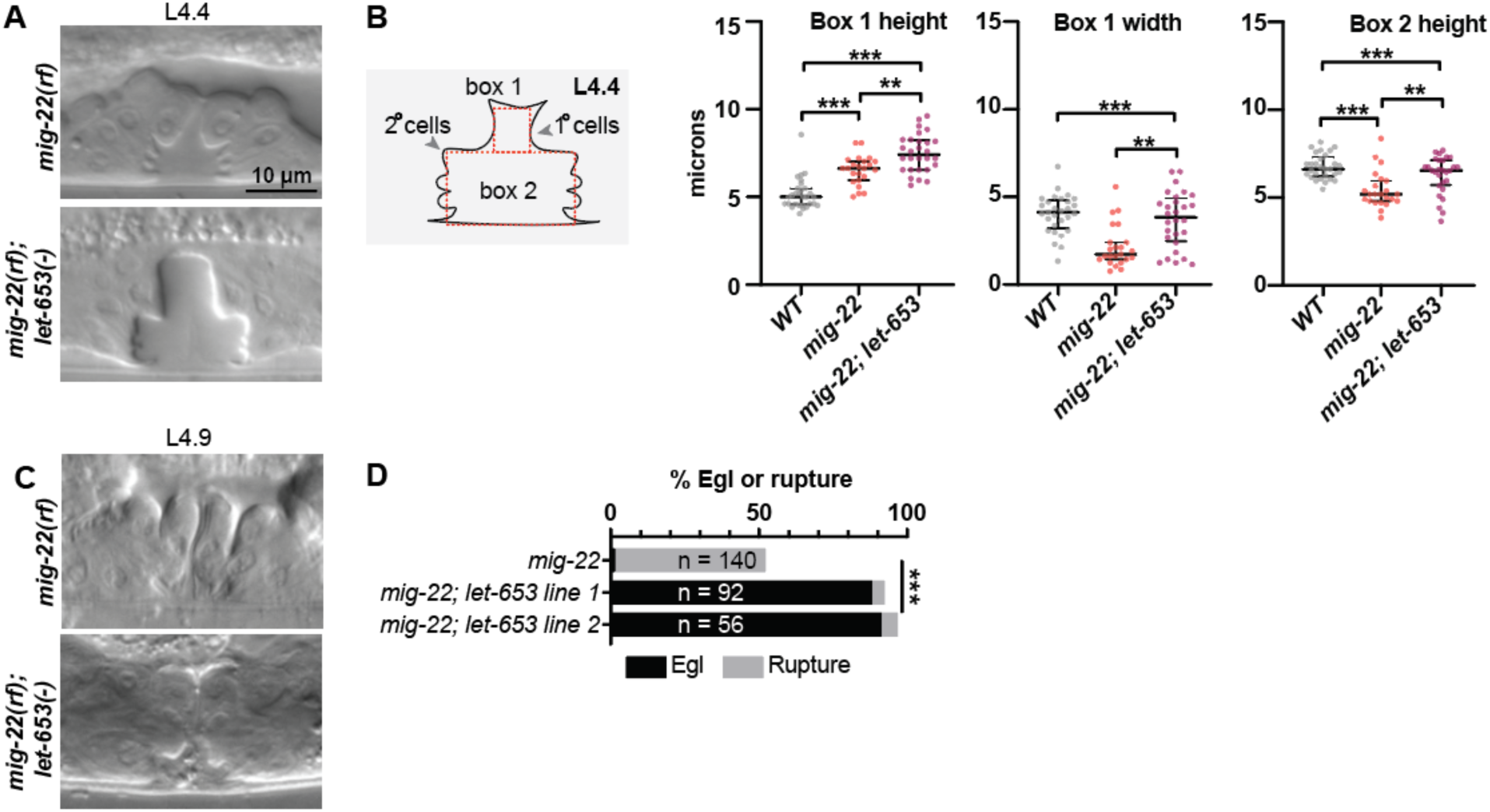
Chondroitin and LET-653 have both lumen-expanding and lumen-constraining roles. A) Loss of *let-653* suppressed the *mig-22(rf)* Sqv phenotype and caused over-inflation of the dorsal lumen. Alleles used were *let-653(cs178)* and *mig-22(k141rf).* B) Lumen dimensions at the L4.4 stage were quantified as in Figure 7C. p values derived from Mann-Whitney U test; **p<0.01, ***p<0.0001. Box 1 height, *mig-22* vs. *mig-22; let-653* p=0.002. Box 1 width, *mig-22* vs. *mig-22; let-653* p=0.001. Box 2 height, *mig-22* vs. *mig-22; let-653* p=0.003. C) At late L4 stages, *mig-22(rf); let-653* double mutants had disorganized material within the vulva lumen (n=10). D) Two independently generated *mig-22(rf); let-653* double mutant strains were analyzed, and both were severely egg-laying defective. ***p<0.0001, Fisher’s exact test.

*mig-22(rf*) mutants still had core fibrils marked by LET-653::SfGFP, though these were narrower and taller than in wild-type, matching the changed dimensions of the lumen (Figure 10D). *mig-22(rf)* mutants also still recruited LET-653(ZP), NOAH-1 and LPR-3 to appropriate apical surfaces (Figure 10A,E). Thus, chondroitin does not appear essential for assembling a sheath matrix per se, though it remains possible that it influences sheath structure in a more subtle way.

### Chondroitin and the sheath matrix have both lumen expanding and lumen constraining properties

To ask if chondroitin and sheath factors work cooperatively to shape the vulva, we examined double mutants between *mig-22(rf)* and *let-653*. Surprisingly, loss of *let-653* largely suppressed the *mig-22(rf)* mutant Sqv phenotype. At the L4.4 stage, double mutants appeared properly inflated in the ventral region, and actually overly inflated in the dorsal, 1°-derived region (Figure 11A,B). Nevertheless, at later eversion stages, the vulva lumen appeared variably abnormal and contained disorganized matrix material (Figure 11C), and as adults, almost all double mutants were Egl or ruptured at the vulva (Figure 11D). These results indicate that MIG-22 and LET-653 have opposing roles in promoting vs. restraining initial lumen inflation, but also have cooperative roles in restraining the subsequent expansion of the dorsal-most 1° vulva toroids, and in promoting later steps of eversion and cuticle formation. These results are inconsistent with the model that chondroitin acts only to exert a uniform hydrostatic expansion force. Rather, as shown here, chondroitin proteoglycans act within a complex luminal scaffold that likely exerts, resists, and distributes multiple different types of vulva cell- and lumen-shaping forces.

## Discussion

The diameter of a tube lumen is ultimately determined by the shape and organization of the cells that surround that lumen. Two well-known determinants of cell shape are the cytoskeleton and the ECM. Here, we showed that the luminal aECM within the developing *C. elegans* vulva tube has a structural and functional complexity that rivals that of the cytoskeleton. The cytoskeleton consists of multiple dynamic and interacting components (actin, microtubules, and intermediate filaments) that are organized into both cytosolic and membrane-anchored fibrils and webs (Fletcher & Mullins, 2010). These cytoskeletal elements can exert both pushing and pulling forces on cell membranes (Fletcher & Mullins, 2010). Similarly, the vulva luminal matrix contains a variety of both free and membrane-attached structural elements. These elements are cell-type specific and highly dynamic over the course of tube morphogenesis. Removal of individual aECM elements, or sets of elements, has distinct effects on cell and lumen shape, revealing both lumen expanding and lumen constricting roles. Ultimately, these data reveal a complex and dynamic aECM which offers a powerful model for investigating aECM assembly, remodeling, and tube-shaping capacity.

### Multiple types of matrix shape the vulva lumen

Although it has been clear for over a decade that chondroitin GAGs are required to inflate the vulva lumen (Herman et al., 1999; Hwang & Horvitz, 2002a, 2002b; Hwang, Olson, Brown, et al., 2003; Hwang, Olson, Esko, et al., 2003), other aECM factors involved in shaping had not been previously described. Here, we showed that the vulva aECM contains multiple discrete elements (Figure 12). First, a granular matrix containing the CPG FBN-1 (and likely many others) fills the luminal cavity by mid-L4. Second, distinct types of membrane-anchored aECMs line different vulva cell types at different stages; these aECM layers contain ZP domain, eLRRon and lipocalin proteins and appear to be analogous to the embryonic sheath that lines the epidermis (Mancuso et al., 2012; Priess & Hirsh, 1986; Vuong-Brender et al., 2017). Third, luminal fibrils form a stalk-like core structure within the central lumen, and then attach to different sheath-covered cell surfaces via ventrolateral fibrils. Each of these elements may play different roles in lumen shaping (Figure 12).

**Figure 12.**
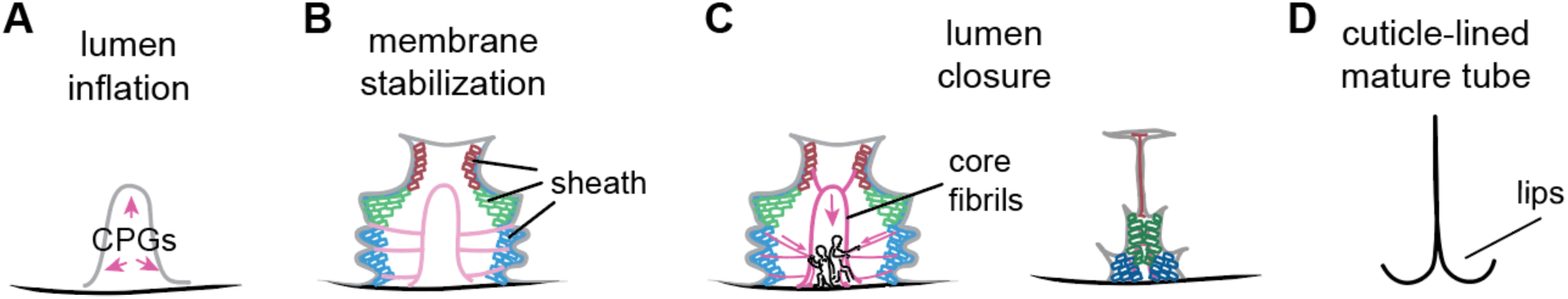
Model for aECM-dependent shaping of the vulva lumen during morphogenesis. A) The vulva lumen is initially expanded by chondroitin proteoglycans (pink arrows). Black; cuticle. Gray; apical membrane. B) A membrane-anchored sheath-like matrix appears alongside matrix fibrils and a central stalk to halt and/or stabilize vulva expansion. vulE/F sheath-like matrix; red. vulC/D sheath-like matrix; green. vulA/B sheath-like matrix; blue. C) The lumen narrows in the anterior-posterior axis. We propose that the central stalk and fibrils attach to the membrane-anchored matrix and pull ventrally, anteriorly, and posteriorly to shape the vulva lumen. The aECM changes over time; sheath components turn over and the membrane-anchored matrix develops cuticle-like features. D) By adulthood, the lumen narrows into a slit and is lined by cuticle.

### Apical matrix may generate, counteract and distribute multiple forces during lumen shaping

Prior work showed that the vulva is expanded by a combination of pushing forces by chondroitin GAGs (Hwang, Olson, Esko, et al., 2003) and actin-myosin constriction by 2° cells, which may then focus the GAG-dependent forces dorsally (Yang, Roiz, Mereu, Daube, & Hajnal, 2017). The data here suggest that GAGs not only inflate the lumen, but also play other roles. First, chondroitin depletion does not affect all vulva cells in the same way; in *mig-22(rf)* mutants, some cells have expanded apical domains, while others have shortened apical domains. Second, in the context of *let-653* loss, *mig-22(rf)* actually causes increased dorsal lumen expansion, revealing a lumen-constraining role for both LET-653 and chondroitin (Figure 11). Finally, the predominant defect in *fbn-1* single mutants is not in lumen inflation, but instead in vulva eversion, showing that GAG-modified proteins have diverse functions in vulva shaping.

The other aECM factors described here may counteract and distribute the forces exerted by GAGs and the cytoskeleton, or may themselves generate additional types of forces. The vulE/F sheath and central core component LET-653 restrains CPG-dependent lumen inflation in early-to-mid-L4 stages, but also helps maintain lumen inflation during the subsequent steps of vulva morphogenesis. Interestingly, LET-653 impacts these later steps even though the sheath-forming LET-653(ZP) domain is no longer present at those stages. LET-653(ZP) and other sheath components may form a membrane-anchored scaffold whose assembly at mid-L4 initially “locks in” a particular lumen size to prevent further CPG-dependent inflation (Figure 12). This scaffold could then serve as a template for recruitment and assembly of later matrix factors, including cuticle collagens, that will sculpt the final structure. Through unknown mechanical connections with membranes and hemidesmosome-like structures, sheath factors also may influence the cytoskeletal organization of vulva cells to promote their rearrangements and cell shape changes. Connections between sheath and muscle at such hemidesmosomes may also serve as anchor points to transmit muscle-generated forces (Vuong-Brender et al., 2017). Finally, the LET-653-marked luminal fibrils appear to constrict during lumen narrowing, and such fibrillar re-organization could potentially exert a pulling force on the sheath and apical membranes (Figure 12), as has been proposed for alae (cuticle ridge) formation (Sapio, Hilliard, Cermola, Favre, & Bazzicalupo, 2005).

Multiple sheath factors likely cooperate in vulva lumen shaping with LET-653. Most sheath factor single mutants, including *let-653*, have only subtle vulva shaping defects and a low percentage of egg-laying defective adults, despite drastic defects in matrix organization (Figures 8, 9). This is in contrast to the much more penetrant and dramatic phenotypes observed in these same mutants in shaping narrow excretory system tubes (Gill et al., 2016; Mancuso et al., 2012) or the embryonic epidermis (Vuong-Brender et al., 2017). The vulva appears to be less sensitive to defects in sheath aECM than these other tissues, possibly because its large lumen can tolerate many irregularities in cell shape and still remain passable for egg-laying.

### LET-653 and cell type-specific partners may traffic through large secretory vesicles in vulF

Individual sheath components appear and disappear from individual cell surfaces at specific timepoints. These localization patterns suggest careful regulation of the sheath aECM. How are these proteins, and their corresponding aECM layers, built and broken down at the correct times and places? Part of the answer may lie in time- and cell-type-specific secretion.

Many secreted matrix proteins are packaged and trafficked via specialized vesicles. For example, collagens are secreted from extra-large vesicles (Malhotra & Erlmann, 2015) and then processed after secretion to enable their assembly (Holmes, Lu, Starborg, & Kadler, 2018). Different ZP proteins secreted from the same cell are sorted into separate pools of vesicles, possibly to prevent their premature association (Jovine, Qi, Williams, Litscher, & Wassarman, 2007). In fact, we observed large vesicles emptying material into the vulva lumen from vulF (Figure 4). LET-653 is a likely cargo within these secretory vesicles and may traffic with one or more partners that form the membrane-anchored sheath matrix over vulE/F. In the absence of LET-653, these partners appear to aggregate abnormally as soon as the vesicle contents are exposed to the luminal environment. Although LET-653 is expressed by all vulva cell types, it may traffic in different types of vesicles or with different partners in each cell type, explaining why LET-653(ZP) does not form the same type of matrix over all vulva cells, and why abnormal vesicles were not observed in other vulva cells in *let-653* mutants.

The vulF secretory vesicles resemble mucin vesicles found in goblet cells of the mammalian lung and gut. Tightly compressed mucin packets in those vesicles are thought to expand rapidly upon reaching the luminal environment due to changes in pH and salt conditions (Birchenough et al., 2015); the CFTR ion channel, which is mutated in cystic fibrosis patients, is expressed on adjacent cell types (Kreda, Davis, & Rose, 2012) and is important for establishing the proper luminal environment for mucin expansion to occur (Kreda et al., 2012). Given that vulF secretory vesicles empty into narrow lumen compartments between vulF and vulE, it is possible that vulE expresses key ion channels or other matrix factors that are important for proper assembly of the extruded matrix.

### The vulva as a system for visualizing and dissecting aECM trafficking and assembly

Classic TEM studies in many systems have shown that aECMs are layered structures (Chappell et al., 2009; Johansson et al., 2011). However, a major challenge for understanding aECMs has been the difficulty in visualizing them in a more high-throughput manner. The large size of the vulva lumen, combined with the transparency of *C. elegans*, allowed us to visualize the various aECM elements by light microscopy in a way that is unprecedented in other systems. The vulva therefore provides a very powerful system for addressing further mechanistic questions regarding how aECM components traffic to the apical surface, how aECM structures are assembled and remodeled, and how they ultimately impact cell and tube shape.

## Materials and Methods

### Worm strains, alleles and transgenes

All strains were derived from Bristol N2 and were grown at 20°C under standard conditions (Brenner, 1974). *let-653* and *let-4* mutants were obtained from mothers rescued with wild-type transgenes expressed in the excretory system under the control of the *lin-48* or *grl-2* promoters (Forman-Rubinsky et al., 2017). *sqv-5* mutants were obtained from heterozygous mothers. All other mutants were obtained from homozygous mothers.

Mutants used included: *lin-12(n137n720)* III (Sternberg & Horvitz, 1989), *lin-12(n137)* III (Greenwald et al., 1983), *mig-22(k141) III* (Suzuki et al., 2006), *sqv-5(n3611) I* (Hwang, Olson, Esko, et al., 2003), *fbn-1(tm290) III* (Kelley et al., 2015), *let-4(mn105) X* (Meneely & Herman, 1979), *sym-1(mn601) X* (Niwa et al., 2009) and *let-653(cs178) IV* (an early nonsense allele) (Gill et al., 2016). See Table S1 for a complete list of all strains used.

Transgenes used included: *csIs64[let-653pro::LET-653::SfGFP; lin-48pro::mRFP] (Gill et al., 2016), csIs66[let-653pro::SfGFP::LET-653(ZP); let-653pro::PH::mCherry; lin-48pro::mRFP] X* (Cohen, Flatt, Schroeder, & Sundaram, 2019), *csEx766[lin-48pro::LET-653::SfGFP; myo-2pro::GFP]* (Forman-Rubinsky et al., 2017), *csEx819[grl-2pro::LET-4; myo-2p::mRFP]* (Forman-Rubinsky et al., 2017), *muIs28[mig-2pro::MIG-2::GFP; unc-31+]* (Honigberg & Kenyon, 2000), *mfIs4[egl-17pro::YFP; daf-6pro::CFP; unc-119+]* (Félix, 2007), *upIs1 [mup-4pro::MUP-4::GFP, rol-6(su1006)] (Hong et al., 2001).* Transgene *aaaIs12 [fbn-1pro::FBN-1::mCherry; ttx-3pro::GFP]* was derived from array *aaaEx78 (Katz et al., 2018)* and expresses full-length FBN-1 tagged internally with mCherry inserted just prior to the ZP domain. *aaaEx78* rescued *fbn-1(tm290)* molt defects and other larval lethality from 85% (n=236) in non-transgenic siblings to 10% (n=1676) in transgenic animals.

CRISPR fusions used included: *let-653(cs262[LET-653::SfGFP]) IV* (this study), *let-4(cs265[mCherry::LET-4]) X* (this study), *noah-1(mc68[NOAH-1::mCherry]) I* (Vuong-Brender et al., 2017), *sym-1(mc85[SYM-1::GFP]) X* (Vuong-Brender et al., 2017), *lpr-3(cs250[SfGFP::LPR-3]) X* (this study), *vab-10(cas602[VAB-10a::GFP])* (Y. Yang et al., 2017).

### Staging and microscopy

Larvae were staged by vulva morphology. Fluorescent, Brightfield, Differential interference contrast (DIC), and Dodt (an imaging technique that simulates DIC) (Dodt & Zieglgansberger, 1990) images were captured on a compound Zeiss Axioskop fitted with a Leica DFC360 FX camera or with a Leica TCS SP8 confocal microscope (Leica, Wetzlar Germany). Images were processed and merged using ImageJ.

For TEM, L4 hermaphrodites from N2 (wild-type) or UP3342 (*let-653(cs178); csEx766[lin-48pro::LET-653::SfGFP; myo-2pro::GFP]*) strains were fixed by high pressure freezing followed by freeze substitution into osmium tetroxide in acetone (Weimer, 2006), and then rinsed and embedded into LX112 resin and cut into serial thin sections of approximately 70 nm each. Sections were observed on a JEM-1010 transmission electron microscope (Jeol, Peabody Massachusetts). Images were processed in ImageJ and manually pseudo-colored in Adobe Illustrator (Adobe, San Jose California).

### CRISPR/Cas9-mediated generation of reporters

To generate LET-653::SfGFP and SfGFP::LPR-3 fusions via CRISPR, a self-excising cassette (SEC) vector containing SfGFP was generated. SfGFP was amplified and inserted a larger fragment of the SEC vector pDD282 (Dickinson, Ward, Reiner, & Goldstein, 2013) via PCR sewing. The large fragment was inserted into pDD282 as a NaeI – BglII fragment. Henceforth, CRISPR was carried out as described (Dickinson, Pani, Heppert, Higgins, & Goldstein, 2015). Briefly, LET-653 or LPR-3 homology arms were inserted into the resulting plasmid and the relevant PAM sites were then mutated via site-directed mutagenesis. sgRNAs were generated via site-directed mutagenesis that inserted a primer encoding the gRNA before a U6 promoter in the plasmid pDD162 (Dickinson et al., 2013). These plasmids were injected into N2 hermaphrodites. F2 progeny were screened by microscopy for Hygromycin resistance and/or SfGFP fluorescence. Insertions were verified by PCR and Sanger sequencing. SfGFP was inserted immediately before the LET-653 stop codon or immediately following the LPR-3 signal peptide. An SEC inserted into an intron within the SfGFP coding sequence was removed via heat shock (Dickinson et al., 2015). Excision was confirmed by PCR and Sanger sequencing.

The mCherry::LET-4 fusion was generated using the SapTrap method (Schwartz & Jorgensen, 2016). LET-4 homology arms were inserted into plasmid pMLS279 using SapI digestion and ligation. The resulting plasmid was co-injected into N2 hermaphrodites with a plasmid containing relevant sgRNAs inserted into pDD162 (Dickinson et al., 2013). F2 progeny were screened by microscopy for mCherry fluorescence. Insertions were verified by PCR and Sanger sequencing. mCherry was inserted immediately after the LET-4 signal sequence.

All CRISPR fusion strains were evaluated for viability. *let-653(cs262[LET-653::SfGFP])* was 93% viable (n=195), *lpr-3(cs250[SfGFP::LPR-3])* was 98% viable (n=137), and *let-4(cs265[mCherry::LET-4])* was 100% viable (n=154).

### Lumen measurements

Vulva dimensions were measured using the box tool in ImageJ. All measurements were performed by a researcher blinded to genotype. Boxes were drawn within the lumen in DIC images and their length and width were recorded. A minimum of 10 animals were examined for each experiment. Statistics were calculated using Prism software (Graphpad, San Diego California) using two-tailed Mann-Whitney U tests.

### Data availability statement

Strains used are listed in Table S1. All strains are available upon request.

## Acknowledgements

We thank Ken Nguyen (Albert Einstein School of Medicine) and Biao Zuo (UPenn Electron Microscopy Resource Lab) for assistance with TEM, Andrea Stout (UPenn CDB Microscopy Core) for training and assistance with confocal imaging, Lily Zekavat for assistance with *lin-12* experiments, and John Murray, Nick Serra and Susanna Birnbaum for helpful discussions and comments on the manuscript. Some strains were provided by the CGC, which is funded by the NIH Office of Research Infrastructure Programs (P40 OD010440). This work was funded by NIH grants R01GM58540 and R01GM125959 to M.V.S., T32 GM008216 and T32 AR007465 to J.D.C., and NIH OD010943 to D.H.H, and by ACS grant RSG-12-149-01-DDC to A.R.F.

## Author Contributions

JDC conceived of and performed experiments, arranged figures, and wrote the paper.

APS performed TEM experiments and interpreted data.

ACB performed experiments.

RFR performed experiments.

DHH offered critical advice and facilities to perform experiments and interpreted data.

HMMN performed experiments and interpreted data.

ARF conceived of experiments and interpreted data.

MVS conceived of experiments, interpreted data, arranged figures and wrote the paper.

