## Supplemental Figures and Table for "A multi-layered and dynamic apical extracellular matrix shapes the vulva lumen in *Caenorhabditis elegans*"

**Figure S1. Ultrastructural features of the mid-L4 vulva aECM**

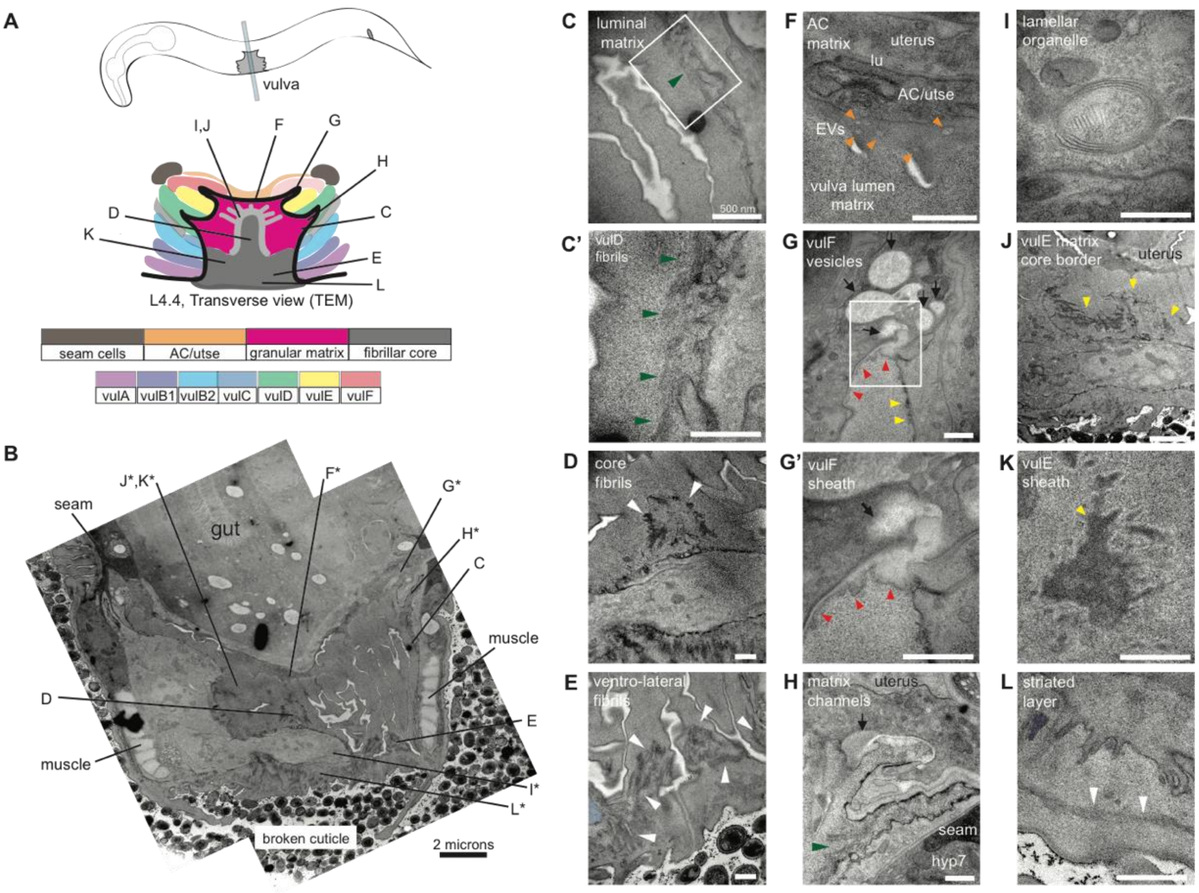


Uncolored images from Figure 4 are shown.

**Figure S2. Ultrastructural features of the late L4 vulva aECM**

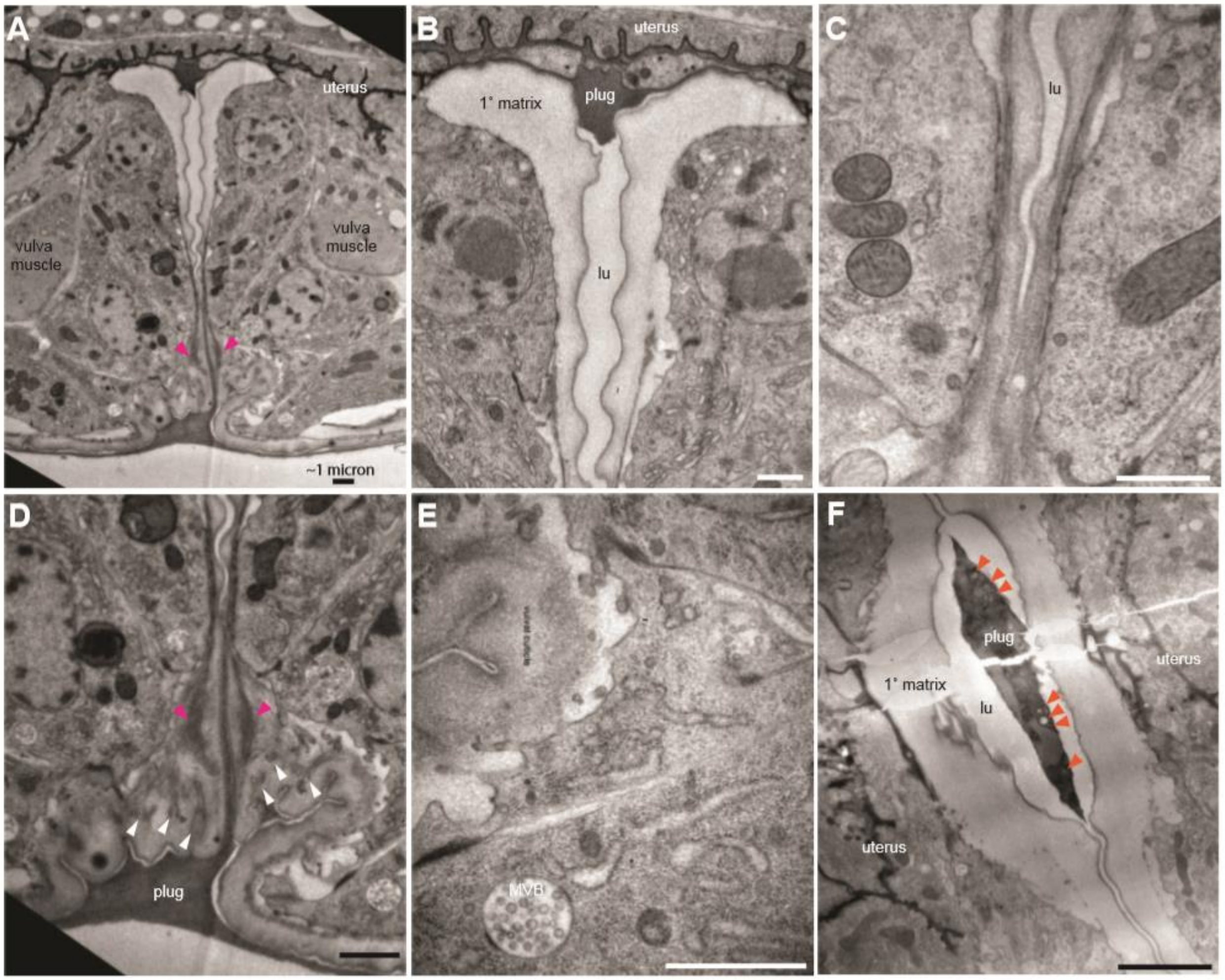


Uncolored images from Figure 5 are shown.

**Figure S3. Measurements of *let-653, mig-22,* and LET-653+ vulvas**

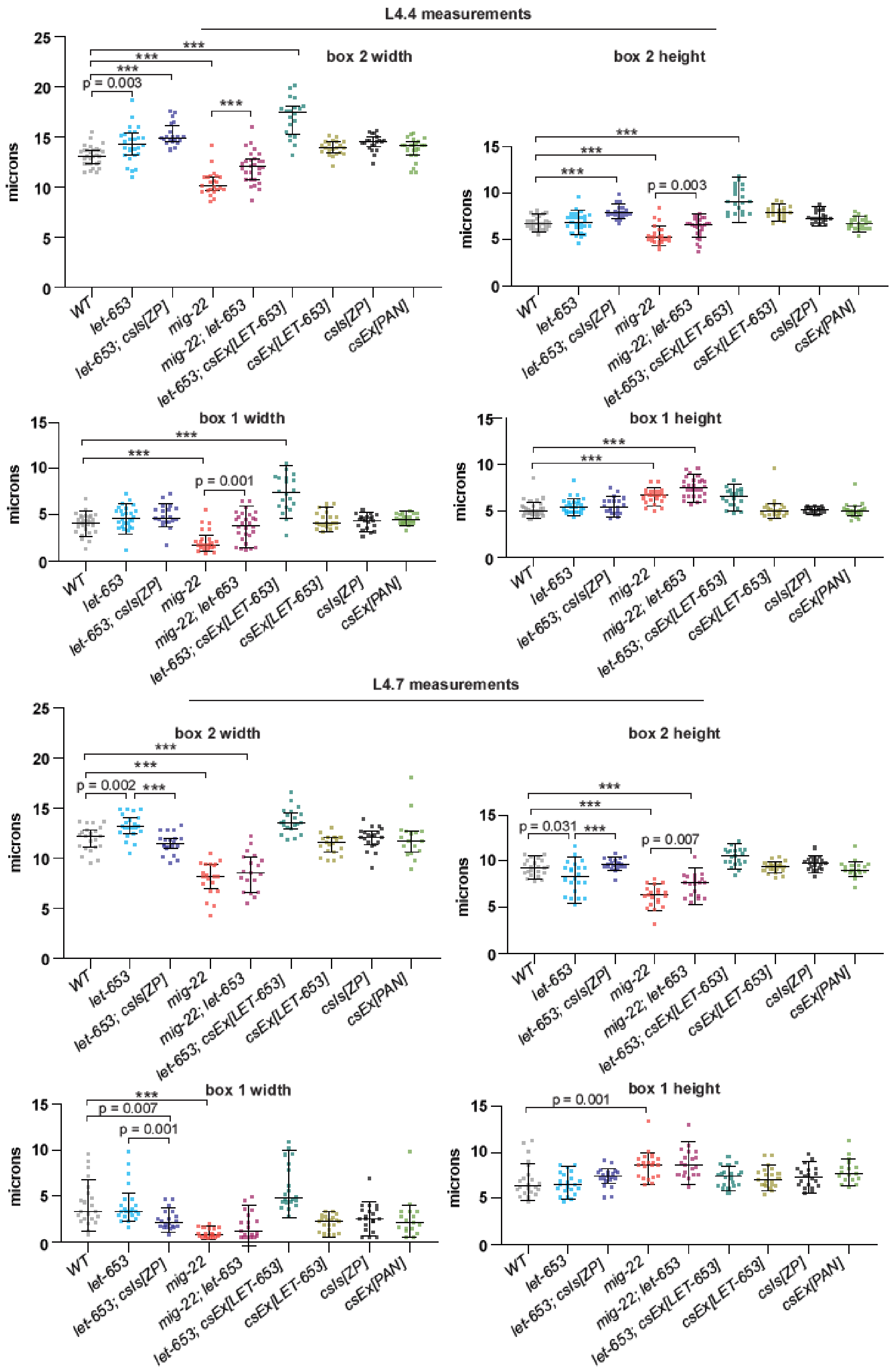

Lumen dimensions at the L4.4 and L4.7 stages were quantified as in Figure 7C. ***$p<0.0001$, Mann-Whitney U test. All p values between 0.01 and 0.0001 are reported. p values above 0.01 are not indicated.

**Table S1. Strains used in this work**

| Strain | Genotype |
|---|---|
| ARF335 | *fbn-1(tm290) III; aaaEx78 [fbn-1pro::FBN-1::mCherry; ttx-3pro::GFP]* (Katz et al., 2018) |
| ARF379 | *aaaIs12 [fbn-1pro::FBN-1::mCherry; ttx-3pro::GFP] V* (Katz et al., 2018) |
| ARF359 | *upIs1 [MUP-4::GFP; rol-6(su1006)] V; aaaEx78 [fbn-1pro::FBN-1::mCherry; ttx-3pro::GFP]* (Hong et al., 2001; Katz et al., 2018) |
| HM24 | *upIs1 V; aaaIs12 V* (Hong et al., 2001; Katz et al., 2018) |
| GOU2043 | *vab-10(cas602 [vab-10a::gfp]) I* (Y. Yang et al., 2017) |
| JU486 | *mfIs4 [egl-17pro::YFP; daf-6pro::CFP; unc-119(+)]* (Mok et al., 2015) |
| ML2482 | *noah-1(mc68 [NOAH-1::mCH(int)]) I* (Vuong-Brender et al., 2017) |
| ML2547 | *sym-1(mc85 [SYM-1::GFP]) X* (Vuong-Brender et al., 2017) |
| NF68 | *mig-22(k141) III* (Suzuki et al., 2006) |
| UP3349 | *aaaIs12 V; csIs64 [let-653pro::LET-653b::SfGFP; rol-6(su1006)]* |
| UP3353 | *let-653(cs178) IV; aaaIs12 V; csEx766 [lin-48pro::LET-653b::SfGFP; myo-2p::GFP]* |
| UP3422 | *csIs66 [let-653pro::LET-653(ZP)::SfGFP; let-653pro::PH::mCherry] X* |
| UP3444 | *sqv-5(n3611)/hT2g I; csIs66 X* |
| UP3666 | *lpr-3(cs250 [ssSfGFP::LPR-3]) X* |
| UP3693 | *noah-1(mc68 [NOAH-1::mCH(int)]) I; lpr-3(cs250 [ssSfGFP::LPR-3]) X* |
| UP3746 | *let-653(cs262 [LET-653::SfGFP]) IV* |
| UP3756 | *let-4(cs265 [ssmCherry::LET-4]) X* |
| UP3757 | *dpy-19(e1259) lin-12(n137)/ hT2g III; csIs66 X* |
| UP3758 | *unc-32(e189) lin-12(n137 n720)/hT2g III; csIs66 X* |
| UP3788 | *noah-1(mc68 [NOAH-1::mCH(int)]) I; let-653(cs262 [LET-653::SfGFP]) IV* |
| UP3856 | *let-653(cs262 [LET-653::SfGFP]) IV; let-4(cs265 [ssmCherry::LET-4]) X* |
| UP3861 | *muIs27 [MIG-2::GFP; dpy-20+]; let-4(cs265 [ssmCherry::LET-4]) X* (Honigberg & Kenyon, 2000) |
| UP3939 | *let-4(mn105) X; csEx819 [grl-2pro::LET-4+; myo-2pro::mRFP]* (Forman-Rubinsky et al., 2017) |
| UP3967 | *mig-22(k141) III; let-653(cs178) IV; csEx766* |
| UP3968 | *mig-22(k141) III; let-653(cs178) IV; csEx766* |
| UP3970 | *mig-22(k141) III; csIs66 X* |
| UP3995 | *muIs28 [MIG-2::GFP; unc-31+]* (Honigberg & Kenyon, 2000) |
| UP3966 | *noah-1(mc68 [NOAH-1::mCH(int)]) I; let-653(cs178) IV; lpr-3(cs250 [ssSfGFP::LPR-3]) X; csEx766* |
| UP4004 | *mig-22(k141) III; muIs28* |
| UP4005 | *let-653(cs178) IV; muIs28; csEx766* |
| UP4014 | *noah-1(mc68 [NOAH-1::mCH(int)]) I; mig-22(k141) III; lpr-3(cs250 [ssSfGFP::LPR-3]) X* |
| UP4025 | *noah-1(mc68 [NOAH-1::mCH(int)]) I; unc-32(e189) lin-12(n137 n720)/hT2g III* |
| UP4027 | *mig-22(k141) III; let-653(cs262 [LET-653::SfGFP]) IV* |

| | |
|---|---|
| UP4038 | *unc-32(e189) lin-12(n137 n720)/hT2g III; lpr-3(cs250 [ssSfGFP::LPR-3]) X* |
| UP4039 | *mig-22(k141) III; mfIs4* |
| UP4040 | *let-653(cs178) IV; mfIs4; csEx766* |
| UP4042 | *unc-32(e189) lin-12(n137 n720)/hT2g III; let-653(cs262 [LET-653::SfGFP]) IV* |
| UP4043 | *dpy-19(e1259) lin-12(n137)/ hT2g III; lpr-3(cs250 [ssSfGFP::LPR-3]) X* |
| UP4044 | *dpy-19(e1259) lin-12(n137)/ hT2g III; let-653(cs262 [LET-653::SfGFP]) IV* |
| UP4045 | *noah-1(mc68 [NOAH-1::mCH(int)]) I; dpy-19(e1259) lin-12(n137)/ hT2g III* |
| UP4047 | *noah-1(mc68 [NOAH-1::mCH(int)]) I; muIs28* |
